# Impact of inflammatory preconditioning on murine microglial proteome response induced by focal ischemic brain injury

**DOI:** 10.1101/2023.04.13.536755

**Authors:** Dario Lucas Helbing, Fabienne Haas, Emilio Cirri, Norman Rahnis, Therese Thuy Dung Dau, Erika Kelmer Sacramento, Nova Oraha, Leopold Böhm, Helen Morrison, Reinhard Bauer

**Author notes:** Corresponding author: Reinhard Bauer, MD, Institute of Molecular Cell Biology, Jena University Hospital, Friedrich Schiller University Hans-Knöll-Straße 2, D-07745 Jena, Germany. The work was performed at the Institute of Molecular Cell Biology, Jena University Hospital, Friedrich Schiller University, Jena, Germany. Email addresses of all authors: Dario Lucas Helbing Fabienne Haas Emilio Cirri Norman Rahnis Therese Thuy Dung Dau Erika Kelmer Sacramento Nova Oraha Leopold Böhm Helen Morrison Reinhard Bauer.

## Abstract

Preconditioning with LPS induces neuroprotection against subsequent cerebral ischemic injury, mainly involving innate immune pathways. Microglia are CNS-resident immune cells that respond early to danger signals through memory-like differential reprogramming. However, the cell-specific molecular mechanisms underlying preconditioning are not fully understood. To elucidate the distinct molecular mechanisms of preconditioning on microglia, we compared these cell-specific proteomic profiles in response to LPS preconditioning and without preconditioning and subsequent transient focal brain ischemia and reperfusion, – using an established mouse model of transient focal brain ischemia and reperfusion. A proteomic workflow, based on isolated microglia obtained from mouse brains by cell sorting and coupled to mass spectrometry for identification and quantification, was applied. Our data confirm that LPS preconditioning induces marked neuroprotection, as indicated by a significant reduction in brain infarct volume. The established brain cell separation method was suitable for obtaining an enriched microglial cell fraction for valid proteomic analysis. The results show a significant impact of LPS preconditioning on microglial proteome patterns by type I interferons, presumably driven by the interferon cluster regulator proteins Stat1/2.

## Introduction

Cerebral ischemic injury is the second leading cause of death and a major cause of long-term disability, with increasing incidence in the young and aging (Collaborators, 2019; Ekker *et al*, 2019; Virani *et al*, 2021). Whilst therapy of acute ischemic stroke induced by large artery occlusion has evolved significantly in recent years, recognizing the value of thrombolysis and/or thrombectomy in appropriately selected patients (Nogueira *et al*, 2018; Thomalla *et al*, 2018), the benefit of reperfusion therapies is incomplete in about half of said cohort (Chamorro *et al*, 2021). Furthermore, ischemia-reperfusion injury is often iatrogenically induced after life-saving endovascular or cardiac procedures (Bonati *et al*, 2010; Sun *et al*, 2012). To combat predictable ischemic injury, preactivation of transient endogenous protective mechanisms (“classical preconditioning”, PC reviewed in (Gidday, 2006; Kariko *et al*, 2004; Kirino, 2002)) induced by stimulation with low doses of an otherwise deleterious insult, has been shown to be effective in reducing and even preventing sequelae of subsequent injurious ischemia (Dirnagl *et al*, 2003; Stenzel-Poore *et al*, 2007).

PC therefore represents a prophylactic intervention with various modalities – including non-injurious ischemia or hypoxia, hypothermia, pharmacological agents, low doses of endotoxin (lipopolysaccharide, LPS) – by inducing mainly inflammatory responses that can limit tissue damage of subsequent transient ischemia/reperfusion (I/R) (Stetler *et al*, 2014; Stevens *et al*, 2014). Notably, PC induced by immune activators Toll-like receptor (TLR) ligands has shown remarkable efficacy in inducing ischemic tolerance: Systemic administration of several TLR ligands, including LPS, prior to focal cerebral ischemia, profoundly reduces ischemic injury in rodent stroke models (Bordet *et al*, 2000; Leung *et al*, 2012; Rosenzweig *et al*, 2007; Stevens *et al*, 2008; Tasaki *et al*, 1997). However, the complex molecular mechanisms underlying PC are not fully understood.

Given such limited therapeutic options and suboptimal tools for diagnosis and prognosis of IS, new strategies to increase brain cell survival after stroke are essential and best achieved by identifying better targets through deeper mechanistic understanding of IS pathophysiology (Hochrainer & Yang, 2022; Lind *et al*, 2015). High-throughput technologies, such as proteomics, have become crucial in unraveling key interactions between different molecular elements in complex biological contexts, such as IS and PC (Horgan & Kenny, 2011).

Previous studies suggest that PC effects of various origins mainly require innate immune pathways, including Toll-like receptors (TLRs) and type I interferons (IFN) (Stenzel-Poore *et al*., 2007; Stevens *et al*, 2011). Microglia, the CNS-resident neuro-immune cells that express these key innate immune receptors and modulators, respond quite early and extremely sensitively to immune-competent signals such as DAMPs and PAMPs, through memory-like differential epigenetic and immunometabolic reprogramming (Lajqi *et al*, 2019; Lajqi *et al*, 2021; McDonough & Weinstein, 2016). It has been shown that PC-dependent specific gene expression signatures in microglia prompt a functional shift of these mutant cells towards an immunomodulatory or protective phenotype (McDonough *et al*, 2017). However, changes in mRNA levels detected by transcriptomics may not correlate with similar changes in protein levels. As proteins perform most essential cellular functions, the composition of the proteome carries critical information about the state of an organism. The proteome is highly dynamic and adapts to changes in the microenvironment through transcriptional, translational, posttranslational and degradational processes (Hochrainer & Yang, 2022).

We focused on the impact of endogenous ischemic protection induced by LPS-induced PC, known to induce a multifaceted reprogramming of brain tissue at risk, leading to markedly reduced tissue destruction by subsequent IS. Thus, we applied a proteomic analysis based on isolated microglia obtained from mouse brains by cell sorting, coupled to mass spectrometry for identification and quantification. This approach was applied to a mouse model of focal brain ischemia and reperfusion by transient middle cerebral artery occlusion (tMCAO), without or after LPS-induced PC (tMCAO_PC), using various bioinformatic analyses to characterize microglial proteome remodeling in response to tMCAO and tMCAO_PC (Fig. 1). This strategy enabled identification of specific effects of LPS-induced PC on transient brain I/R and potential neuroprotective effects mediated by microglial reprogramming.

**Figure 1:**
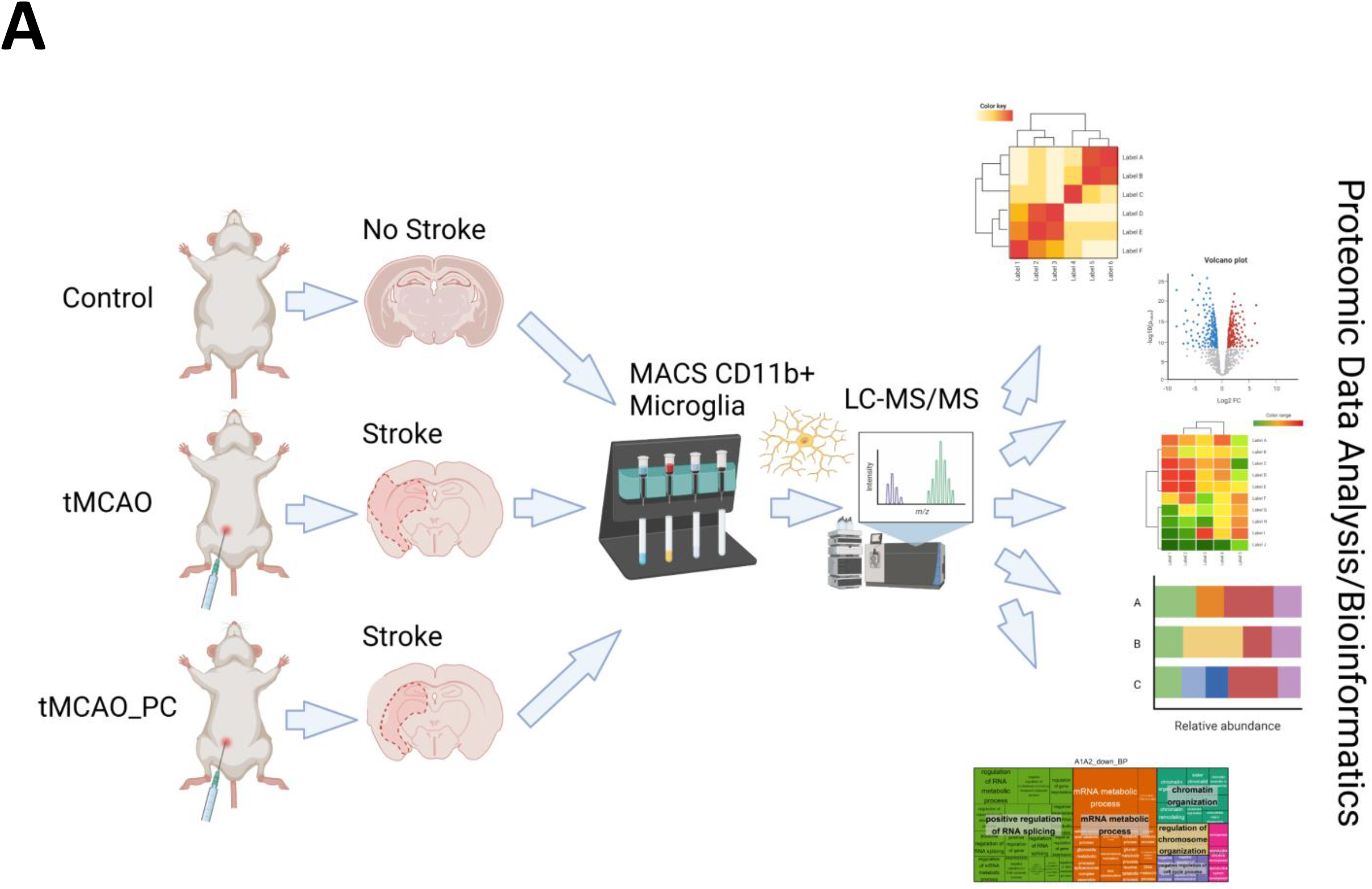
Schematic synopsis of performed experimental strategy to perform proteomic analysis of microglial response on temporal focal brain ischemia and reperfusion, with (tMCAO) or without previous LPS-induced inflammatory preconditioning (tMCAO_PC). (This figure was created under an institute license (Leibniz Institute on Aging Jena) with Bio-Render.com).

Our data confirm that LPS-induced PC caused a significant reduction in infarct volume. The brain cell separation method used is suited to obtaining an enriched microglial cell fraction for valid proteomic analysis. The results show a significant impact of type I IFN on brain I/R-induced and LPS-induced PC modified microglial proteome patterns, presumably driven by cluster 3 regulators Stat1/2.

## Results

### LPS preconditioning reduced cerebral infarct size following I/R

LPS-induced preconditioning led to an approximate halving of the infarct volume (Fig. 2A). In addition, preconditioned animals displayed significantly better neurological recovery within the first 48h after tMCAO, as determined by modified Bederson score (Bederson *et al*, 1986; Schmidt *et al*, 2016) (Fig. 2B). LPS-induced preconditioning only slightly worsened the clinical course prior to tMCAO (Table EV1).

**Figure 2:**
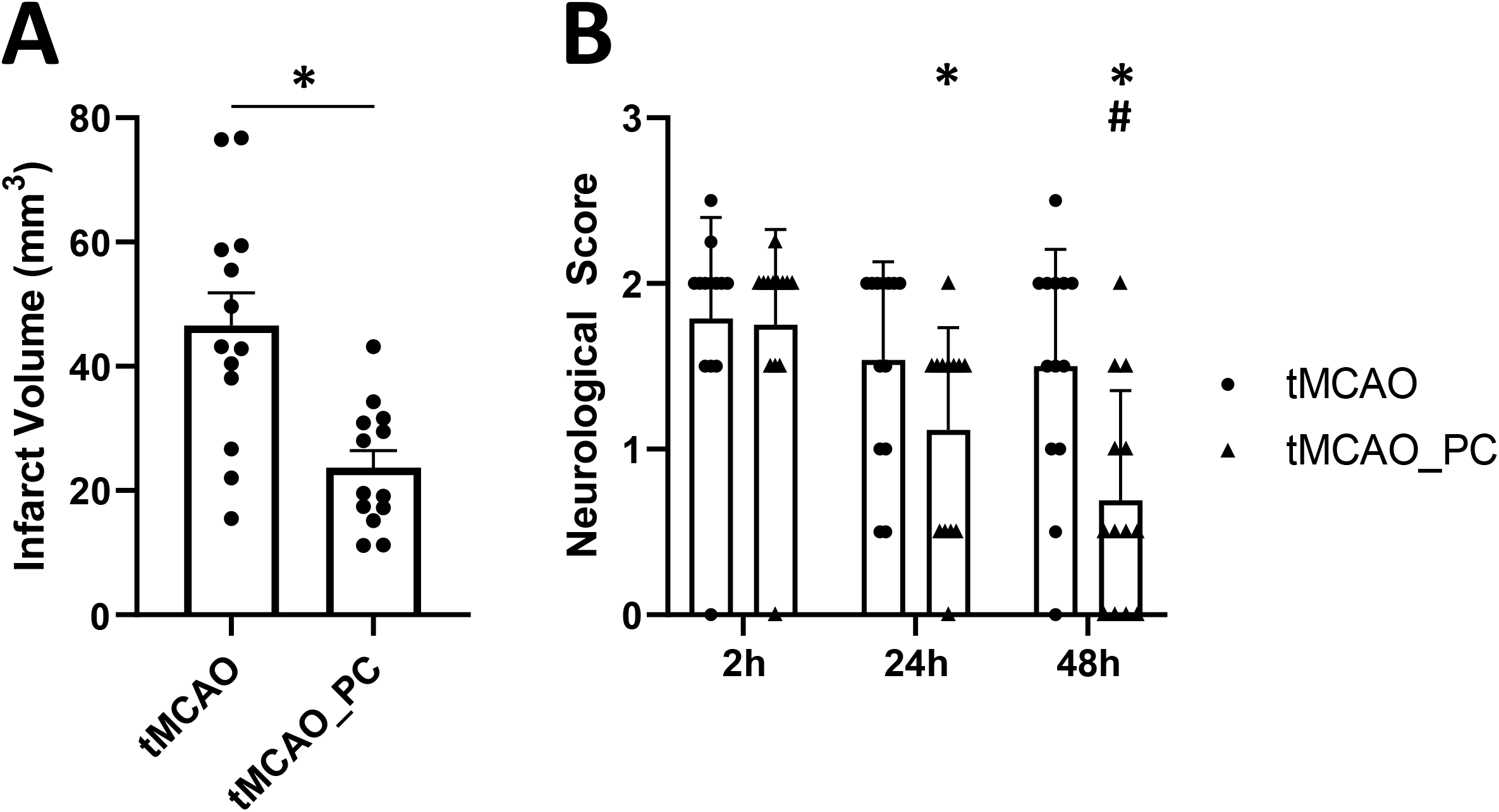
Reduction in infarct size and improved neurological recovery in LPS-preconditioned animals and tMCAO. **A** Reduced infarct volume in animals with LPS-induced preconditioning (n = 13 animals per condition, scatterplot, mean + SEM, statistical power: 0.99, effect size (Cohen’s d): 1.61, Unpaired Students t-test with Welch’s correction with * = p < 0.05, specifically p = 0.0012). **B** Time course of neurological deficits after tMCAO as detected by a modified Bederson score (Bederson *et al*., 1986; Schmidt *et al*., 2016) shows reduction in symptom severity within 48 hours post-tMCAO, with significantly better neurological recovery in LPS-preconditioned animals (n = 13 animals per condition, two-way ANOVA with p = 0.0002 for the factor time post-tMCAO and p = 0.025 for the factor preconditioning. Post-hoc Holm-Sidak test with * p < 0.05 for the comparison to 2h timepoint within each conditioning group (tMCAO or tMCAO_PC), p = 0.0084 for the comparison 2h vs. 24h tMCAO_PC, p < 0.0001 for the comparison 2h vs. 48h tMCAO_PC, p = 0.0508 for the comparison 48h vs. 24h in the tMCAO_PC group, ^#^p < 0.05 for comparisons tMCAO vs. tMCAO_PC within each timepoint, p = 0.0050 for the 48h timepoint.)

### MACS-based microglia isolation

To ensure high-quality proteomic profiles of isolated microglia with response to focal brain I/R and preconditioning, the magnetic-activated cell sorting-based isolation protocol for adult microglia was validated by flow cytometric data analysis (Fig. EV 1A) and proteomic analyses (Fig. EV 1B-E) of microglia cells (CD11b+), as well as the CD11b negative cell fraction (CD11b-/NTCF (= non-target cell fraction)) comprised of astrocytes, oligodendrocytes and neurons. Principal component analysis of the proteomic profiles of both fractions revealed marked differences, proving the presence of different cell types (Fig. EV 1B). Integration of the proteome profiles determined here with previously determined proteome profiles of different cell types from adult mouse brain (Sharma *et al*, 2015), revealed a distinct presence of microglia specific marker proteins in the CD11b+ fraction; non-microglia proteins were almost completely absent, whereas non-microglia proteins were mainly found in the CD11b- fraction (NTCF) (Fig. EV 1C, D). The same pattern was observed by investigating commonly used marker proteins for microglia (Aif1/Iba1), astrocytes (Gfap), oligodendrocytes (Cnp) and neurons (Tuj1/Tubb3) (Fig. EV 1E) – indicating a comparatively good enrichment of microglia by MACS-based separation and sparse contamination of non-microglial cells. Therefore, the isolation method used is suitable for obtaining an enriched microglia cell fraction for valid proteomics experiments.

### Differential proteomic response in microglia after tMCAO with or without LPS preconditioning

To categorize global proteomic changes in microglia induced by focal brain injury and inflammatory preconditioning, a principal component analysis was performed on the total proteomes of individual microglial cells isolated from three experimental cohorts (naïve microglia, microglia from brains after tMCAO, microglia from brains after tMCAO with previously induced LPS preconditioning). A clear separation and clustering by tMCAO alone, as well as by preconditioning and tMCAO, was found (Fig. 3A). Furthermore, all tMCAO samples were separated along the first principal component (explaining ∼ 37.4% of the total variance) from untreated non-tMCAO samples (CTRL), indicating that tMCAO in general induces similar characteristic changes in microglia, independent of preconditioning. Nevertheless, 11.5% of sample variation is explained by the second principal component, wherein a clear separation is visible between preconditioned tMCAO (tMCAO_PC) and non-preconditioned tMCAO microglia (tMCAO); suggesting specific additional changes within microglial proteome induced by pre-conditioning. As such, we performed a differential abundance analysis to identify the proteomic changes commonly induced by tMCAO and by tMCAO with preconditioning (tMCAO versus CTRL and tMCAO_PC versus CTRL, abbreviated with A1 and A2, respectively, Fig. 3B). We also performed comparative analyses of up- or downregulated proteins in A1 and A2 to specifically identify the differentially regulated proteins induced by preconditioning. While the majority of up- or downregulated proteins were shared (414 upregulated, 498 downregulated) between the A1- and A2- comparisons, a substantial amount were significantly altered when affected mice received LPS preconditioning before tMCAO (243 upregulated, 309 down- regulated, Fig. 3C). This is also reflected in the correlation plot (Fig. 3D), where a strong correlation between changes in A1 and A2 was identified both for the whole proteome (Spearman’s ρ = 0.83) and for proteins identified as significantly changed in both comparisons (q-value ≤ 0.05, Spearman’s ρ = 0.91). Therefore, the generalized proteomic response of non-preconditioned and preconditioned microglia to tMCAO is generally similar (Fig. 3D). Among the significantly up- and downregulated proteins are several candidate proteins associated with inflammation, as well as restriction of inflammation. Notably, TLR2, a critical receptor for the microglial response to ischemia (Tang *et al*, 2007), was among the top 10 upregulated proteins in comparisons A1 and A2 (Fig. 3D). Several other interferon related proteins (e.g., Stat1, Ifit1) appeared to be upregulated, particularly in preconditioned microglia (Fig. 3D). Moreover, the lysosomal protein CD68 (Kurtin & Bonin, 1994) was more upregulated after preconditioning (log2 fold change (Log2FC) A2 = 2.57 vs. Log2FC A1 = 1.27). Amongst the most downregulated genes in preconditioned microglia was Mef2c, a factor downregulated in microglia due to the presence of Type I IFN, e.g., during aging (Deczkowska *et al*, 2017) (Log2FC A2 = −3.11, Fig. 3D), but was not identified in non-preconditioned microglia. A large number of proteins related to splicing and histones were also downregulated in both comparisons (Fig. 3D).

**Figure 3:**
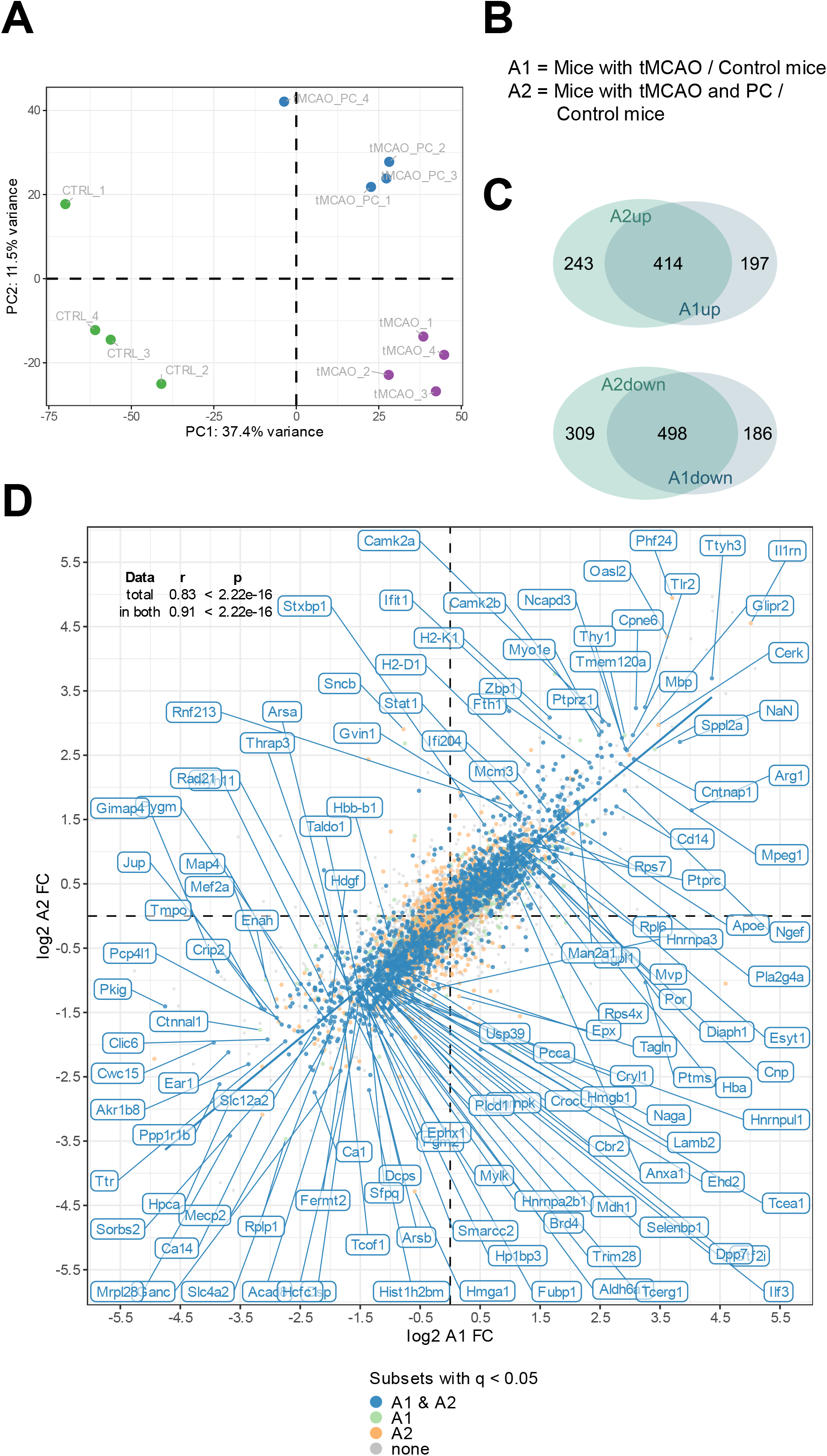
Global microglial proteome changes in response to tMCAO and tMCAO with preconditioning. **A** Principal component analysis of the total proteome of non-preconditioned microglia, 2 days after tMCAO (A1) and LPS-preconditioned microglia 2 days after tMCAO (A2) (n=4, each). All tMCAO and tMCAO_PC samples are separated via the first principal component from the control microglia, indicating strong proteome changes by tMCAO in microglia. In addition, tMCAO_PC samples are separated from tMCAO microglia by PC2, indicating preconditioning- specific proteome changes after tMCAO in microglia with good reproducibility. **B** Definition of the relevant comparisons for subsequent differential abundance and enrichment analyses. **C** Venn diagrams illustrating the number of proteins being significantly (│log2 FC│≥ 0.58, meaning 50% change and q-value ≤ 0.05) either up- (A1up, A2up, upper panel) or downregulated (A1down, A2down, lower panel), either in both datasets (overlapping area) or uniquely in one of both datasets. **D** Correlation plot of the log2 FCs of the A1 and A2 comparison. In the upper left corner, the correlation statistics for total as well as significant proteomes are depicted. (As indicated, in both A1 and A2 significantly changed proteins are labelled in blue, in A1 significantly changed proteins are labelled in light green, in A2 significantly changed proteins are labelled in orange and non-significant proteins are colored in gray. DEPs are labelled with their respective name, if for one of both comparisons │log2 FC│≥ 2.7 and the q-value ≤ 0.05, or if │log2 FC│≥ 1 and q-value ≤ 10^-9^; Log2FC: log2 fold change).

### KEGG pathway and gene ontology enrichment analysis revealed changes in immune response, metabolism, debris clearance and transcriptional/translational processes

The protein lists derived from Venn diagrams were subjected to gene ontology enrichment and KEGG pathway enrichment analysis using DAVID (Huang da *et al*, 2009), to identify and differentiate specific biological processes that are either generally changed by tMCAO, or only changed by preconditioning.

Amongst the most prominent regulated KEGG pathways were terms related to mRNA processing. Specifically, the “spliceosome” and “RNA degradation” pathways were enriched in the protein lists of downregulated proteins in both non-preconditioned and preconditioned microglia (A1 and A2, respectively) – with more proteins significantly associated with both terms in preconditioned microglia (56 proteins in A2 versus 45 proteins in A1), indicating potentially stronger downregulation (Fig. 4). The “mRNA surveillance” KEGG pathway was uniquely enriched in the list of downregulated proteins in preconditioned microglia, suggesting a possible alteration of alternative splicing and RNA processing events upon LPS preconditioning. Furthermore, proteins belonging to the ribosomal machinery and DNA replication were upregulated in A1 and A2, alluding to microglial activation associated with cytokine production and secretion after focal ischemia. The KEGG terms “protein export” and “protein processing in endoplasmic reticulum” were also upregulated in both conditions, reinforcing the notion that ribosomal proteins are upregulated, as cytokine production is induced in response to tMCAO and cytokines are often post-translationally modified prior to export and secretion (Vanheule *et al*, 2018) (Fig. 4). As expected, the KEGG pathway “phagosome” was strongly enriched in the list of upregulated proteins of microglia under both conditions, reflecting the phagocytic activity of microglia after ischemic stroke (Schilling *et al*, 2005) (Fig. 4). Slightly more phagosome DEPs were found in preconditioned microglia (26 vs. 22 in non-preconditioned microglia), whilst 9 phagosome DEPs were found to be uniquely upregulated in preconditioned microglia. Of interest, several synaptic proteins were highly upregulated in preconditioned microglia, represented by the KEGG term “synaptic vesicle cycle”, indicating neuron/synapse phagocytosis of activated microglia (Fig. 4); the Pathview visualization (Luo & Brouwer, 2013) shows an apparent induction of phagocytosis-associated proteins at all levels - uptake, maturation and processing - in both microglia sets, with several proteins either only or more strongly upregulated in preconditioned microglia (Fig. EV 2A). We also retrieved a list of proteins annotated to be localized to phagocytic vesicles (GO:0045335 “Phagocytic vesicle”) as a proxy for phagosome content. Heatmap visualization and quantification of protein intensities of all mapped phagocytic vesicle proteins, showed that the majority of phagocytic vesicle proteins were upregulated after tMCAO in both microglia sets, with more proteins induced in preconditioned microglia (Fig. EV 2B, C).

**Figure 4:**
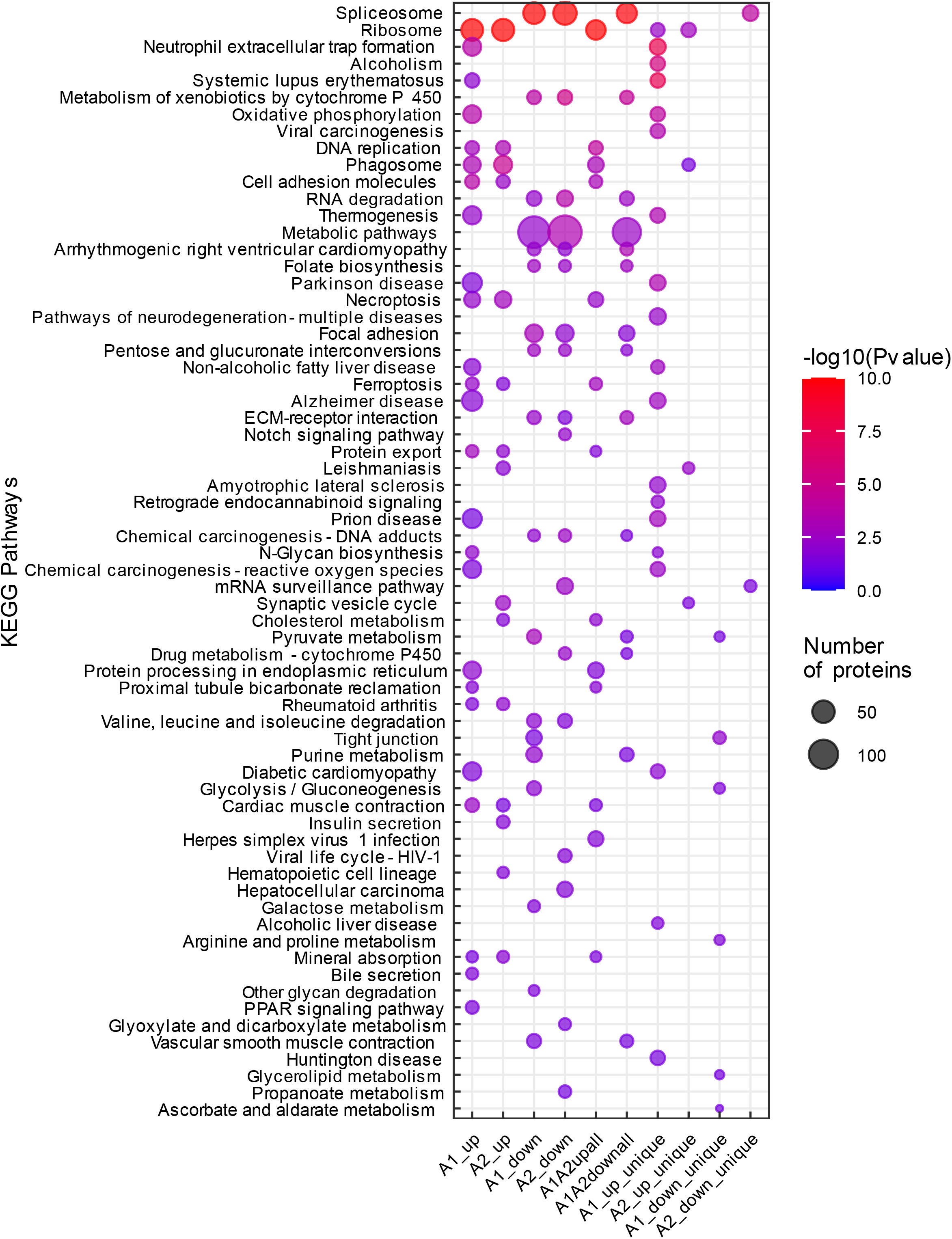
KEGG pathway enrichment analysis. **A** Functional pathway analysis of DEPs with KEGG pathway enrichment analysis. Analyses were performed on proteomic data obtained from non-preconditioned microglia, 2 days after tMCAO (A1) and LPS-preconditioned microglia 2 days after tMCAO (A2). All DEPs (│log2 FC│≥ 0.58 and q-value ≤ 0.05) were submitted to the DAVID platform for KEGG pathway enrichment analysis. The subsequent list was filtered for pathways with a Fisher’s Exact p-value ≤ 0.05, which were considered to be significantly enriched. (As indicated, circle size represents the number of proteins associated with the respective enriched pathway, colour scale specifies the level of significance. The experimental groups A1/A2_up/down, denoting the up-and downregulated groups from the single comparisons tMCAO vs. control and tMCAO_PC vs. control, and thereafter the protein lists derived from the Venn diagrams, are depicted on the x-axis.

The “oxidative phosphorylation” KEGG pathway was uniquely enriched in non-preconditioned microglia. (Fig. 4). However, Pathview and heatmap visualizations showed the majority of oxidative phosphorylation associated proteins were also upregulated in preconditioned microglia, at lower levels (Fig. EV 3A, B); confirmed by quantification of protein intensities of all mapped oxidative phosphorylation proteins (Fig. EV 3C). Although iNOS was not detected in this proteomics experiment, NADPH oxidase Nox2 – the other major ROS producing enzyme in microglia – was markedly upregulated in non-preconditioned microglia (Log2FC A1 = 1.76, q-value = 0.0009), whereas it was about 50% less upregulated in preconditioned microglia (Log2FC A2 = 0.85, q-value = 0.0009) (Fig. EV 3D). Overall, the KEGG pathway enrichment analyses performed indicate biological changes in the microglial response to tMCAO and preconditioning generally similar, but subsequently “fine-tuned”, in terms of greater or lesser induction of key microglial effector functions/phenotypes after tMCAO.

### Gene ontology enrichment (GO) analysis reveals remodeling of microglial inflammatory and functional response to tMCAO with preconditioning

While KEGG pathways often contain biologically simplified pathways, GO-terms can provide more detailed biological insights by describing molecular functions, biological processes or cellular components associated with enriched proteins (Harris *et al*, 2004). Using the DAVID platform (Huang da *et al*., 2009), additional enrichment analyses for level 5 biological process and cellular component gene ontology terms were conducted. The enrichment results were subjected to semantic similarity analysis, clustering and visual dimensionality reduction analysis (Gu & Hubschmann, 2022). GO enrichment analyses for biological processes revealed two large GO-clusters containing metabolic and RNA/nucleotide and cytoskeleton related changes, primarily found in the lists of downregulated proteins (Fig. 5), concordant to the KEGG pathway enrichment related changes (Fig. 4). Concomitantly, cellular component GO-terms, enriched in the list of downregulated proteins, were nearly all related to the spliceosome, chromatin and cytoskeletal components (Fig. EV 4). Several enriched GO-term clusters were detected in different lists of upregulated proteins related to inflammatory activation, vesicle transport pathways such as phagocytosis and endocytosis, and oxidative stress (Fig. 5). Correspondingly, upregulated proteins mapped or were enriched in cellular component GO-terms related to the respiratory chain and mitochondrial proteins, in addition to vesicles such as the endosome and neuronal and synapse-associated proteins – all indicative of phagocytosis of neuronal debris by microglia (Fig. EV 4).

**Figure 5:**
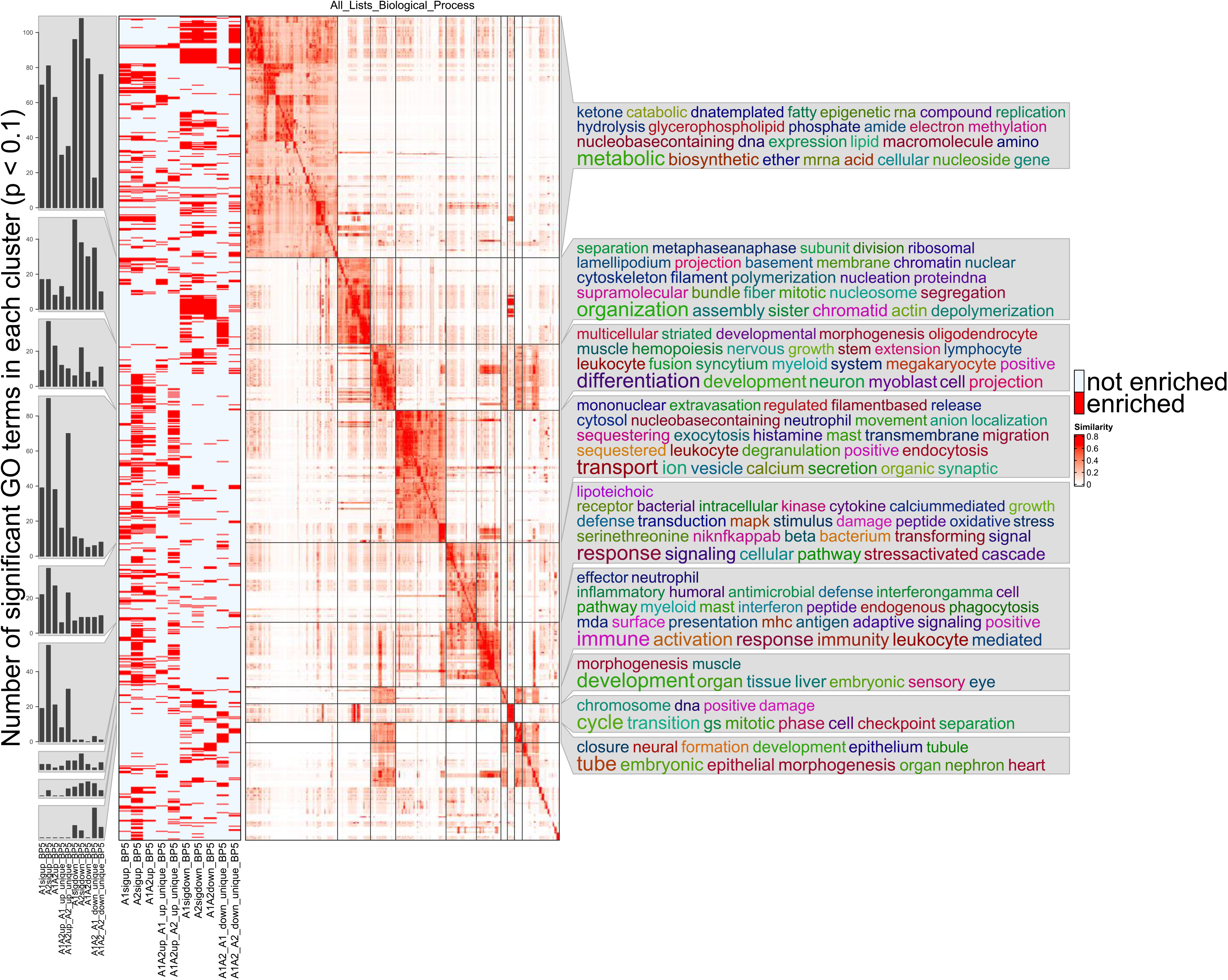
Gene ontology enrichment analysis for biological processes. Functional gene ontology enrichment analysis of level 5 GO-terms of biological processes. Analyses were conducted based on proteomic data obtained from non-preconditioned microglia, 2 days after tMCAO (A1) and LPS- preconditioned microglia, 2 days after tMCAO (A2) the experimental groups A1/A2_up/down, denoting the up-and downregulated groups from the single comparisons tMCAO vs. CTRL and tMCAO_PC vs. CTRL, and thereafter the protein lists derived from the Venn diagrams. Enriched GO-terms with a Fisher’s Exact p-value ≤ 0.1 were used for semantic similarity clustering – the similarity matrix with specific clusters is depicted in the middle. Each row corresponds to one GO-term; in the matrix each column corresponds to one GO-term. On the left panel a binary heatmap is displayed showing whether a GO-term was enriched in the protein target list or not, where each row corresponds to one GO-term. On the extreme left bar plots are shown, where cluster-wise the number of GO-terms for each protein list is counted in a cluster-wise manner. On the right a word cloud is annotated to the similarity matrix, providing information about the biological processes in keywords, in terms of an overrepresentation of individual words from all GO-terms located in this cluster.

Notably, for almost all generally induced GO-term clusters functionally related with tMCAO, more GO-terms were always found to be enriched in the in A2 upregulated protein list, or uniquely present in the A2 upregulated protein list. The opposite can be observed for proteins associated with oxidative stress. These data suggest a fine-tuning of the microglial response by enhancing or attenuating the upregulation of proteins induced by tMCAO in general (A1) (Bar plots of the corresponding clusters, Figure 4 and S2 (new S4)).

To investigate the upregulated GO-terms indicating biological process as well as cellular component in more detail, the different semantic similarity clusters were plotted in treemaps. Among the different clusters related to inflammatory activation, a specific cluster of GO-terms enriched in the list of proteins upregulated in A1 and A2 was related to cytokine responses: (i)“response to interferon-alpha”, (ii) “response to interferon-beta”, (iii) “response to type I interferon” and in general (iv)“cellular response to cytokine stimulus”, signifying a potential importance of type I IFN (Interferon-alpha and Interferon-beta (Ivashkiv & Donlin, 2014)) in modifying microglial activation after tMCAO (Fig. EV 5A).

Furthermore, cellular component GO-terms enriched in upregulated proteins of non-preconditioned (A1) and preconditioned (A2) microglia were related to mitochondrial proteins, vesicle proteins (especially phagocytic and endocytic vesicle membranes), ribosomal proteins and axonal/synaptic proteins (Fig. EV 5B). Plotting the GO-terms enriched uniquely in upregulated proteins of preconditioned microglia, largely resembles the pattern of the enriched GO-terms of both A1 and A2 upregulated proteins (Fig. EV 6A, B), supporting the assumption that LPS-induced preconditioning modifies ischemia-dependent proteome response of microglia and related biological effector functions.

### Preconditioning alters I/R-associated microglial inflammatory phenotype via Type I IFN signaling

The conducted KEGG pathway enrichment and gene ontology enrichment analyses indicated substantial inflammatory activation of microglia after tMCAO, irrespective of precedent treatment (Fig. 5). Considering that microglia can adopt pro- and anti-inflammatory phenotypes after I/R, proteomes form LPS- non-preconditioned and preconditioned microglia were compared with the proteomic core signatures of pro- and anti-inflammatory microglia activation states, as determined previously (Bell-Temin *et al*, 2015). The detected overlap of proteins revealed a profound induction of a large set of pro- and anti-inflammatory signature proteins significantly changed in tMCAO and tMCAO_PC microglia (Fig. EV 7A, B). Quantification of the pattern of upregulated DEPs overlapping to either pro- or anti-inflammatory signature proteins, unveiled markedly higher protein intensities of both signatures owing to tMCAO, but increased upregulation after preconditioning (Fig. EV 7A, B). Specifically, the heatmap of pro-inflammatory marker proteins shows three separated clusters: (i) a small cluster of proteins downregulated in microglia with tMCAO, (ii) a larger cluster of proteins quite similarly upregulated in tMCAO and tMCAO_PC microglia, (iii) an additional cluster of proteins of markedly stronger upregulation in preconditioned microglia (Fig. EV 7A, B). Proteins belonging to the latter were Ifih1, Gvin1, Rnf213, Bst2, Ifit1, Parp9, Ifit3, Gbp2, Cmpk2, Isg15, Pyhin1, Marcksl1, Tap2 and Sqstm1.

A heatmap of anti-inflammatory proteins with similar clustering, showed the third cluster of proteins was markedly larger compared to that found in the heatmap of proinflammatory proteins (Fig. EV 7A). Proteins belonging to that cluster were Pyhin1, Irgm1, Ifit3, Iigp1, Gvin1, Oasl2, Ifit1, Parp9, Ifih1, Gbp2, Cmpk2, Isg15, Parp14, Dhx58, Trex1, Stx18 and Sqstm1. These proteins belong to clusters of proteins more strongly induced in preconditioned microglia and are either part of the cellular type I IFN response or modulate type I IFN signaling. Hence, data support the assumption that inflammatory activation and induction of microglial pro- and anti-inflammatory signature proteins are likely driven and characterized by type I IFN induction, which appears to be enhanced in LPS-preconditioned microglia.

A gene set enrichment analysis (GSEA) was performed on both proteomic datasets to provide further evidence, using a RNAseq analysis dataset obtained from adult microglia after *in vivo* IFN-β treatment (Deczkowska *et al*., 2017). Those genes upregulated by IFN-β were used to perform a gene set enrichment analysis (GSEA) with our two datasets. Significant enrichment of the type I IFN microglia signature was observed in tMCAO-microglia proteomes and a slightly stronger enrichment in the preconditioned microglia. Heatmap visualization and quantification of the protein intensities of IFN-β dependent proteins showed that the majority is more strongly upregulated in the preconditioned microglia (Fig. 6A-C) –data that substantiate a significant impact of type I IFN on tMCAO induced and by LPS preconditioning modified microglial proteome patterns.

**Figure 6:**
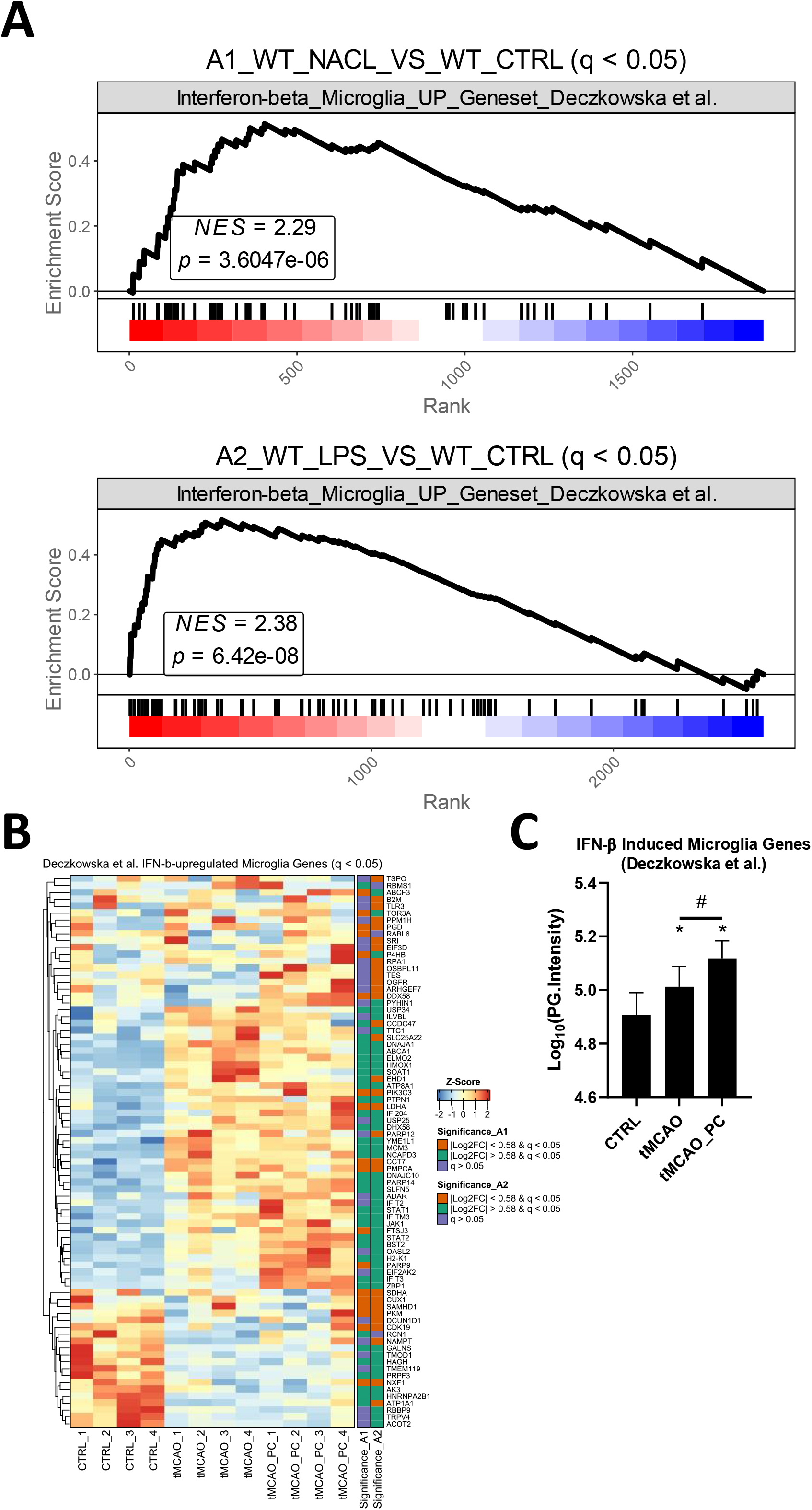
tMCAO-related microglial inflammatory activation resembles a type I IFN-signature. **A** Gene set enrichment analyses for all by IFN-b in microglia induced genes (Geneset from(Deczkowska *et al*., 2017)) Upper plot: Visualization and statistics of the enrichment of microglia interferon signature in the A1 comparison. Lower plot: Visualization and statistics of the enrichment of microglia interferon signature in the A2 comparison. **B** Heatmap visualization of the z-score normalized protein intensities of all samples Only proteins are shown that were significantly changed in A1 or A2 (q ≤ 0.05) and belong to the IFN-β induced gene signature (Deczkowska *et al*., 2017) (n=4, each). On the right side of the heatmap an annotation is shown as to whether a protein was significantly changed in A1 or A2 and if yes, above or below the fold change cut-off of │log2 FC│≥ 0.58. **C** Bar charts of Log10(PG.Intensities) of all in A1 or A2 significantly changed proteins belonging to the IFN-β induced gene signature (n=80 proteins, a Friedman test was conducted, with a p-value = 0.0001 with Post-hoc Wilcoxon signed-rank tests with * = p < 0.05 for comparisons to the control group, specifically p = 0.0022 for CTRL vs. tMCAO, p = 0.0003 for CTRL vs. tMCAO_PC and # = p < 0.05 for the comparison tMCAO to tMCAO_PC, specifically p = 0.0026). All p-values were adjusted for multiple testing with Bonferroni-Holm correction.

Type I IFN signaling is highly diverse, involving multiple pathways and co-regulated protein networks designed to provide robust host defense without inducing autoimmunity (Ivashkiv & Donlin, 2014). Meanwhile, a detailed description of different evolutionarily conserved interferon network clusters has been published (Mostafavi *et al*, 2016). These contain both regulator proteins and target proteins (i.e., interferon signature genes (ISG)) and were obtained and mapped to the tMCAO-associated microglia proteome data. Visualization of the mapped network clusters shows a distinct enrichment of cluster 3 (C3) (Fig. 7A). Both regulator and target proteins of cluster 3 are upregulated in tMCAO-associated microglia. In preconditioned microglia, considerably more C3 target proteins were upregulated (13 proteins in A1; 24 proteins in A2) compared to non-preconditioned microglia (Figure 6A). Quantification of all C3 regulator and target protein intensities showed significant upregulation of C3 regulator proteins in preconditioned microglia (Fig. 7B). Furthermore, C3 target proteins increased their protein intensities in non-preconditioned microglia subjected to tMCAO, alongside a markedly increased upregulation in preconditioned microglia (Fig. 7B).

**Figure 7:**
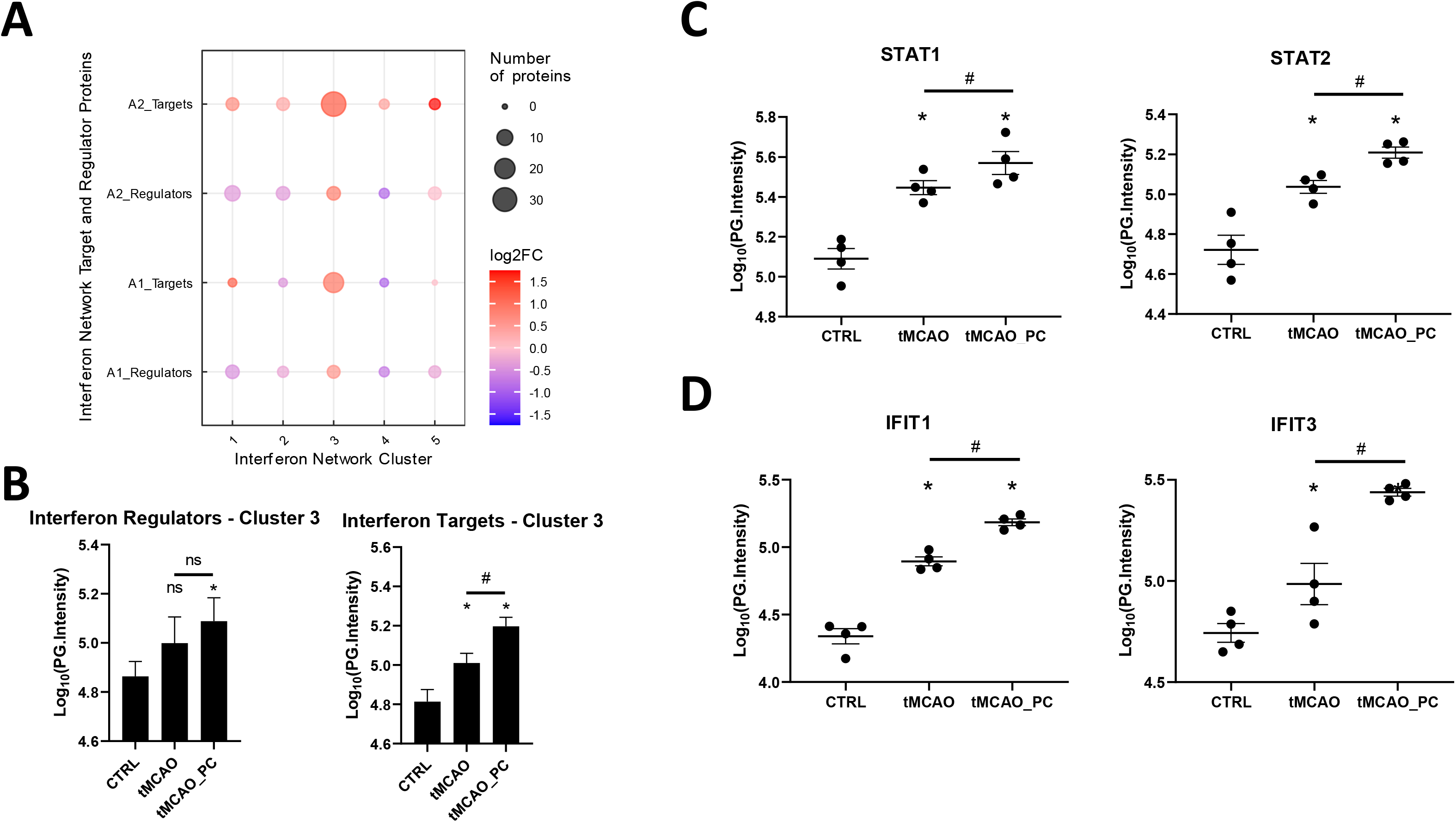
The type I IFN microglial signature after tMCAO is likely driven by Stat1/2. **A** Mapping of significantly changed proteins to the IFN-network of regulators and interferon target genes as determined previously by (Mostafavi *et al*., 2016). **B** Bar charts of Log10(PG.Intensities) of all in A1 or A2 upregulated significantly changed proteins belonging to cluster 3 IFN-regulators (left plot), or cluster 3 IFN-targets (right plot) (n=7 proteins for IFN-regulators, n=32 proteins for IFN-targets. A Repeated measures ANOVA was conducted, with a p-value = 0.0401 for regulator and p < 0.0001 for target proteins. Post-hoc Holm-Sidak tests were conducted with * p < 0.05 for comparisons to the control group and ^#^ p < 0.05 for the comparison tMCAO to tMCAO_PC, as indicated in the respective figure. Specifically, p = 0.0397 for CTRL vs. tMCAO_PC for regulator proteins and for target proteins p < 0.0001 for all 3 comparisons). **C, D** Bar chart scatter plots of Log10(PG.Intensities) of Stat1 and Stat2, belonging to cluster 3 IFN-regulators (**C**) and Ifit1, Ifit3, representing cluster 3 IFN-targets (**D**), as stated previously (Mostafavi *et al*., 2016) (n=4, each). Shown are the FDR-adjusted p-values (=q-values) as reported by Spectronaut software in the differential abundance analysis; *q ≤ 0.05 for comparisons to the control group and ^#^ q ≤ 0.05 for the comparison tMCAO to tMCAO_PC. Specifically, q-values for Stat1 were q = 0.0473 for tMCAO vs. CTRL, q = 8.011E-14 for tMCAO_PC vs. CTRL and q = 0.00016 for tMCAO_PC vs. tMCAO. q-values for Stat2 were q = 0.0289 for tMCAO vs. CTRL, q = 0.0084 for tMCAO_PC vs. CTRL and q = 0.0156 for tMCAO_PC vs. tMCAO. q-values for Ifit1 were q = 0.00056 for tMCAO vs. CTRL, q = 3.7663E-05 for tMCAO_PC vs. CTRL and q = 0.00059 for tMCAO_PC vs. tMCAO. q-values for Ifit3 were q = 0.001 for tMCAO vs. CTRL, q = 5.4312E-05 for tMCAO_PC vs. CTRL and q = 5.3175E-07 for tMCAO_PC vs. tMCAO.

The major C3 regulators were studied to more specifically verify effects of tMCAO and microglial preconditioning on C3 interferon network activit. While Irf9 was not identified, Irf7 was unaltered. In contrast, Stat1 and Stat2, the main constituents of the interferon stimulated gene factor 3 (ISGF3) transcription factor complex (Mostafavi *et al*., 2016), were markedly induced in tMCAO-associated microglia. Intriguingly, a stronger induction in preconditioned microglia was verified (Figure 6C), while two representative members of Interferon-induced protein with tetratricopeptide repeats (IFIT) protein family – Ifit1 and Ifit3 – and thus prominent C3 target proteins, showed increased upregulation in LPS-preconditioned microglia (Fig. 7D). Thus, the significant effect of type I IFN on I/R-induced and LPS-preconditioned microglial proteome patterns is likely driven by the cluster 3 regulators Stat1/2.

## Discussion

Microglia are likely the first responders to ischemia, given their ceaseless surveillance of the microenvironment for alterations (Davalos *et al*, 2005; Nimmerjahn *et al*, 2005), and undergo dose-dependent reprogramming and metabolic shift as a consequence of TLR4 priming (Lajqi *et al*., 2021; McDonough *et al*., 2017; Nair *et al*, 2019). Hence, microglia are viewed as key target cell type for improved insights into the complex molecular reprogramming due to brain I/R and endogenous neuroprotection by PC (McDonough *et al*, 2020). Brain I/R and PC-related changes in microglial gene expression signatures have been widely described (Hamner *et al*, 2022; Hamner *et al*, 2015; McDonough *et al*., 2017; McDonough & Weinstein, 2020), but less is known about the extent and functional impact of proteomic alterations in microglia. For the first time, we comprehensively report the effects of LPS-induced PC on microglial proteomic changes induced by transient MCAO in adult mice.

Our data confirm previous cell-specific and genomic reports that preconditioning, which reduces the deleterious consequences of focal brain I/R, is apparently driven by biological processes associated with modified microglial activation and upregulation of specific inflammatory responses that modulate microglial pro-inflammatory activity (Gesuete *et al*, 2012; McDonough & Weinstein, 2020; Rosenzweig *et al*, 2004). Specifically, an enhanced enrichment of type I IFN in signature was observed in microglial proteomes after preconditioning and subsequent focal brain I/R (Fig. 6). Parsing the evolutionarily conserved interferon network clusters involved (Mostafavi *et al*., 2016), revealed an enhanced upregulation of cluster 3 regulator proteins in preconditioned microglia (Fig. 7). These findings prove on the translational level that the previously suggested type I IFN signaling in microglia does play a critical role in multiple forms of preconditioning-mediated protection (McDonough & Weinstein, 2020). The advantage of proteomics over other high throughput omics techniques is its assessment of the end product arising from a combination of adaptations on every level. This is particularly relevant in stroke, after which translation is stalled and altered mRNA levels, detected by transcriptomics, may not correlate with similar changes in protein levels (DeGracia, 2017; Hochrainer & Yang, 2022). Yet, it must be considered that the effects of type I IFNs may be time-dependent: administration of IFNβ shortly before or after tMCAO exerts a protective effect against ischemic stroke, with a significant reduction in infarct volume through its anti-inflammatory properties targeting reperfusion injury, which is observed after a few days and one or three weeks (Kuo *et al*, 2016; Marsh *et al*, 2009; Veldhuis *et al*, 2003a; Veldhuis *et al*, 2003b). Repeated systemic application of IFNβ for three or seven days failed to protect against experimental ischemic brain injury, determined one week after tMCAO in rats. This was associated with significant weight loss and alterations in hematology and chemistry profiles (Maier *et al*, 2006). Therefore, early temporal activation of type I IFN signaling system yields resistance to subsequent prolonged ischemic exposure. This temporal pattern sits well with the concept of microglial priming (Garcia-Bonilla *et al*, 2014; Lajqi *et al*, 2023 subm; Lajqi *et al*., 2019) and allows the possibility that downstream microglial pathways, including those involving ISG products, are critical effectors in mediating protection and/or enhanced recovery (McDonough & Weinstein, 2020).

To determine the PC-induced modulation of effector mechanisms, or cellular functions, known to be involved in key microglial response patterns to brain I/R, KEGG pathway analyses were performed to examine microglial ROS production and phagocytic activity. Considering that ROS production in microglia underlies three main sources (i) the mitochondrial respiratory chain where ROS accumulate physiologically, (ii) inducible nitric oxide synthase (iNOS) and (iii) cellular NADPH oxidases (NOX), in microglia mainly Nox2 (Simpson & Oliver, 2020; Zhu *et al*, 2022), our data showed a markedly reduced upregulation of microglial Nox2 due to PC, which may be associated with the reduction in infarct size by LPS-PC. These results confirm a previous report which showed that LPS-induced PC is accompanied by a reduced infarct size and preserved neurovascular function (Kunz *et al*, 2007). Phagocytosis is generally considered a beneficial process that leads to clearance of potentially harmful cellular components and may also contribute to the resolution of neuroinflammation after ischemic stroke (Neumann *et al*, 2009; Schmidt *et al*., 2016). It has been shown that upregulation of microglial phagocytic activity occurs rapidly after brain ischemia (Stoll *et al*, 2004) and represents the predominant phagocytic activity (Schilling *et al*., 2005) for removal of released danger signals resulting from dying cells and debris after necrotic cell death. Type I IFNs also act directly on microglia to modulate phagocytosis, including the clearance of degenerating axons, degraded myelin and apoptotic cells (Hosmane *et al*, 2012; Rajbhandari *et al*, 2014), suggesting intimate interplay between LPS- PC-induced modulation of type I IFN signaling and enhanced microglial phagocyte activity.

Still, some limitations must be considered – as we used the entire affected hemisphere for microglial isolation, no differential proteome analysis could be performed for infarct core, penumbra and surrounding brain tissue. Furthermore, we used CD11b for cell-sorting of microglial cells from brain tissue. Even when brains have been cleared of intravasal blood, invading monocytes/macrophages also express the surface integrin CD11b (Bennett *et al*, 2016; El Khoury *et al*, 2007; Hickman *et al*, 2013). However, up to three days after ischemic stroke, invading monocytes/macrophages represent a minority of all CD11b+ cells (Jayaraj *et al*, 2019; Schilling *et al*., 2005; Werner *et al*, 2020). Schilling et al. showed that there were no infiltrating macrophages on the first day and that the number of infiltrating macrophages remained very low on day 2 after 30 min MCAO (Schilling *et al*., 2005). Werner et al. showed by lineage tracing of blood-born macrophages that less than 20% of all Iba1-positive cells represent infiltrating macrophages at day 3 after IS (Werner *et al*., 2020).It can therefore be assumed that the majority of observed proteomic changes documented here can be assigned to microglia. As we only examined the effects of LPS-PC at a time point (48h post-tMCAO) at which is known from previous studies that microglia are the most active immunocompetent cells in ischemic brain tissue, follow-up studies characterizing longer-term effects are required.

In summary, we have provided translational evidence that distinct microglial reprogramming, inducing modified proteomic patterns, underlies endogenous tolerance to transient brain ischemia. Multifunctional adaptations such as inflammatory activation, phagocytosis and ROS production occurred in microglial cells in response to I/R, all of which were differentially modulated by LPS-PC. In particular, LPS-PC was characterized by the induction of an evolutionarily conserved type I interferon response, which may be driven mainly by Stat1/2 activation. Our results offer insights into the regulation of the innate immune response to the ischemic/reperfused brain and immunological preconditioning, which can be starting points for further studies aimed at modulating neuroinflammation to induce neuroprotection due to brain ischemia and reperfusion.

## Materials and Methods

### Animals and experimental procedures

Male 12- to 16-week-old C57BL/6J mice were used in this study. The animal procedures were performed according to the guidelines of Directive 2010/63/EU of the European Parliament on the protection of animals used for scientific purposes. Experiments were approved by the Thuringian State Office for Food Safety and Consumer Protection. Efforts were made to reduce the number of animals used and their suffering. All surgeries were performed under appropriate anesthesia (see below).

Animals were introduced at least one week before commencing the interventions, to ensure appropriate acclimatization (Obernier & Baldwin, 2006) and housed at neutral ambient temperature (30±0.5°C) (Gordon *et al*, 1998) during the entire experimental period.

Following acclimatization, mice were injected randomly with LPS (0.8 µg/g b.w., i.p.) for induction of immunological preconditioning (Dirnagl *et al*, 2009),or physiological saline as a single intraperitoneal injection. Additionally, saline (500 μl) was injected subcutaneously immediately after LPS administration and after 24 h and 48 h. Meloxicam (Metacam®), used to prevent or treat pain, was administered orally 1-2mg/kg after LPS/saline injection, 24h and 48h.

### Focal cerebral ischemia/reperfusion

Three days later animals received transient focal brain ischemia and reperfusion (I/R) by temporal unilateral middle cerebral artery occlusion (tMCAO). Prior to surgery, mice received meloxicam 1-2mg/kg orally. Subsequently, animals were anesthetized with 2.5 % isoflurane for induction and 1.5 % isoflurane for maintenance in 70/30 % nitrous oxide/oxygen, administered by mask. Rectal temperature was maintained at 36.5 to 37 °C with a feedback-controlled heating blanket. Temporal middle cerebral artery occlusion (tMCAO) was induced using the intraluminal filament technique (Huang *et al*, 1994). Briefly, the right common carotid artery, the external carotid artery and the internal carotid artery (ICA) were dissected from surrounding tissue. A 7–0 nylon monofilament (70SPRe, Doccol Corp, USA) was inserted into the ICA (11 mm) to occlude the middle cerebral artery. Operation time per animal did not exceed 15 min. The intraluminal suture was left in situ for 45 min. Animals were re-anesthetized and the occluding monofilament withdrawn to allow reperfusion. Animals were allowed to survive for 48 hours. Two hours post recovery from anesthesia, as well as 24 and 48 hours later, neurological deficits were scored in order to verify correct tMCAO induction by a modified Bederson score (Bederson *et al*., 1986; Schmidt *et al*., 2016) (scoring system: 0, no deficit; 1, forelimb flexion; 2, unidirectional circling; 3, longitudinal spinning; 4, no movement/death). Mice were excluded from analysis when subarachnoid hemorrhage was macroscopically observed during brain harvesting, or died before the end of the allotted reperfusion time. No difference was observed in exclusion rates between the groups. 48 hours after reperfusion, brains were harvested for subsequent immunohistochemical analyses or microglial cell sorting, accordingly.

### Infarct volume estimation

Infarct volume was determined as described previously (Schmidt *et al*., 2016). Mice were deeply anesthetized and perfused with 4 % paraformaldehyde (PFA) in phosphate buffer, after rinsing with PBS by cardiac puncture via the left ventricle. Brains were removed immediately after fixation and postfixed for 5 h in 4 % PFA at 4 °C. After cryoprotection in phosphate-buffered saline (PBS) containing 30 % sucrose, brains were frozen in methylbutane at −30 °C and stored at −80 °C. Whole brains were cut by coronal sections at 40 μm on a freezing microtome (Microm International GmbH, ThermoScientific, Germany). The slices were immunostained by MAP2 (see below) to visualize the infarctions. Sections were photographed with a digital Olympus DP50 camera. Planimetric measurements (ImageJ software, National Institutes of Health, Bethesda, MD) blinded to the treatment groups were used to calculate lesion volumes, which were then corrected for brain edema.

### Magnetic-activated cell sorting (MACS) of primary microglial cells

For microglial cell isolation mice brain vessels were rinsed transcardially by ice-cold PBS perfusion and brains harvested immediately after. The cerebellum and meninges were removed and split in the median sagittal plane. In each case, right hemisphere containing infarction, when mice suffered from MCAO or being unaffected (sham cohort), was sliced into small pieces and transferred to customer-specified tubes for magnetic-activated cell sorting (MACS) (Adult Brain Dissociation Kit #130-107-677 and Cd11b MicroBeads, mouse #130-093-634, Miltenyi Biotec, Bergisch Gladbach Germany). Samples were processed per manufacturer’s instructions with the following modifications: Initially 1% HEPES 1M was added for pH buffering to the D-PBS and PB buffer. Samples were transferred to a C Tube containing the enzyme mix I (enzyme P and buffer Z from the Adult Brain Dissociation Kit #130-107-677, Miltenyi Biotec, mixed according to manufacturer’s instructions). Immediately thereafter, enzyme mix II (enzyme A and buffer Y, from the Adult Brain Dissociation Kit #130-107-677, Miltenyi Biotec, mixed according to manufacturer’s instructions) was added and the samples processed using a gentleMACS Octo Dissociator with Heaters (gentleMACS Program 37C_ABDK_01). The samples were temporarily centrifuged, then triturated 10 times with two different-lumen Pasteur pipettes. Next, the cell suspensions were filtrated with a MACS SmartStrainer and 10 ml of cold D-PBS added, followed by centrifugation for 10 min at 300 g, 4 °C. The supernatant was removed and the cell pellets resuspended in 3100 µl D-PBS. Subsequently, 900 µl of Debris Removal Solution was added and mixed. The mixture was gently overlayed with 4 ml of D-PBS and mixing of the phases was avoided. Three phases formed after a centrifugation step at 3000 g for 10 min, 4 °C. The upper two phases were removed and 15 ml of D-PBS was added to the remaining cell pellet, followed by three inversions of the tube. Following another centrifugation step (1000 g for 10 min, 4 °C), the cell pellet was resuspended in 90 µl PB buffer. 10 µl of CD11b (Microglia) MicroBeads were added and mixed. This mixture was incubated for 15 min at 4 °C in the dark and mixed every 5 min by slightly flicking the tube. The cells were washed by adding 1 ml of PB buffer, centrifuged at 300 g for 5 min, 4 °C, then resuspended in 500 µl of PB buffer.

For flow cytometric analysis, 2x 20 µl samples were collected (unstained original fraction and original fraction (“OF”)). Next, MS Columns were placed onto a MACS® Separator, and prepared by rinsing with 500 µl of PB buffer. The whole brain cell suspension was then applied to the columns and the flow-through containing unlabeled cells collected, hereafter referred to as non-target cell fraction (“NTCF”). 100 µl of cell suspension was collected from this fraction for FACS analysis. The column was washed 3x with 500 µl of PB buffer, removed from the separator and placed into a collection tube. 1 ml of D-PBS was pipetted onto the column and immediately afterwards the magnetically labeled CD11b positive cells were flushed out by pushing the plunger into the column. This step was repeated once. 100 µl cell suspension was collected from this fraction again for FACS analysis, hereafter referred to as “Cd11b+” ≙ CD11b positive fraction. This fraction was centrifuged at 300 g for 5 min, 4 °C, followed by aspiration of the supernatant. The resulting cell pellet was snap-frozen in liquid nitrogen and stored at −80 °C, pending further analysis.

### Flow cytometry and data analysis

Three different cell suspensions were obtained: (i) the original fraction of unstained cells before column separation, (ii) the non-target cell fraction consisting of the flow-through of CD11b negative cells and (iii) the CD11b positive fraction containing microglia. 100 µl of each cell suspension were used (original fraction contains 20 µl of total brain cell suspension with 80 µl of PB buffer added) for analysis of MACS-sorted primary microglial cells and 5 µl of CD11b+-APC antibody added (1:20). These samples were incubated for 10 min in a refrigerator, protected from light. The cell suspensions were then washed by adding 1 ml of PB buffer, followed by centrifugation at 300 g for 5 min, 4 °C. Next, the supernatant was completely aspirated and the cell pellet resuspended in 200 µl of PB buffer. Finally, the cells were analyzed by fluorescence measurement (BD FACSCantoTM System). The collected data were exported for further analysis and generation of scatter plots with FlowJo (version 10.2, BD Bioscience).

### Mass spectrometry

#### Sample preparation for proteomics

Snap frozen cell pellets were resuspended in lysis buffer (final concentration: 0.1 M HEPES/pH 8; 2 % SDS; 0.1 M DTT) and vortexed. All samples were sonicated using a Bioruptor (Diagenode) (10 cycles with 1min on and 30s off with high intensity @ 20 °C). For reduction and full denaturation of the proteins, the lysates were incubated at 95 °C for 10 min, before treatment with iodacetamide (room temperature, in the dark, 30 minutes, 20 mM).Each sample was subsequently treated with 8 volumes of ice-cold acetone to 1 volume sample and left overnight at −20 °C to precipitate the proteins. Next, samples were centrifuged at 14,000 rpm for 30 minutes, at 4 °C. Following removal of the supernatant, the precipitates were washed twice with 200 μL of 80% acetone (ice cold). After each wash the samples were vortexed and centrifuged for 2 minutes at 4°C. The pellets were allowed to air-dry before being dissolved in digestion buffer (1M Guanidine, 100 mM HEPES, pH 8) with sonication (3 cycles in the Bioruptor as above) and incubated for 4 h with LysC (1:100 enzyme: protein ratio) at 37 °C, with shaking at 600 rpm. Next, the samples were diluted 1:1 with deionized water (from a Milli-Q® system) and incubated with trypsin (1:100 enzyme: protein ratio) for 16 h at 37 °C. In the presence of a slow vacuum, the digests were acidified with 10% trifluoroacetic acid and desalted (Waters Oasis® HLB μElution Plate 30μm). The columns were conditioned with 3x100 μL solvent B (80% acetonitrile; 0.05% formic acid) and equilibrated with 3x 100 μL solvent A (0.05% formic acid in milliQ water), before the samples were loaded, washed 3 times with 100 μL solvent A then eluted into PCR tubes with 50 μL solvent B. These eluates were dried down with a speed vacuum centrifuge and dissolved in 50 μL 5% acetonitrile, 95% milliQ water, with 0.1% formic acid, prior to analysis by LCMS/MS.

#### LC-MS data dependent (DDA) and independent (DIA) acquisition

For data dependent acquisition (DDA), peptides were separated using UltiMate 3000 UPLC system (Thermo Fisher Scientific), fitted with a trapping column (Waters nanoEase M/Z Symmetry C18, 5μm, 180 μm x 20 mm) and an analytical column (Waters nanoEase M/Z Peptide C18, 1.7μm, 75μm x 250mm). Solvent A consisted of water, 0.1% formic acid, while solvent B consisted of 80% (v/v) acetonitrile, 0.08% formic acid. The samples (500 ng) were loaded with a constant flow of solvent A at 5 μL/min onto the trapping column, with a trapping time of 6 minutes. Peptides were eluted via the analytical column at a constant flow of 0.3 μL/min. During this step, the percentage of solvent B increased in a linear fashion from 1 % to 7 % within 7 minutes, before further increasing to 32 % within 24 more minutes and finally to 50 % within a further 8 minutes. The outlet of the analytical column was coupled directly to a Q exactive HF (Thermo Fisher Scientific) using the Proxeon nanospray source. The peptides were introduced into the mass spectrometer via a Pico-Tip Emitter 360 μm OD x 20 μm ID; 10 μm tip (New Objective) and a spray voltage of 2.2 kV applied. The capillary temperature was set at 300 °C and the S-lens RF value to 60%. Full scan MS spectra with mass range 350-1650m/z were acquired in the Orbitrap at a resolution of 60,000 FWHM (= full width half maximum). The filling time was set at maximum of 20 ms with an automatic gain control (AGC) target value of 3x10^6^ ions. A Top15 method (selecting the top 15 most intense m/z features) was applied to select precursor ions from the full scan MS for fragmentation (minimum AGC target of 1x 10^3^ ions, normalized collision energy of 31%), quadrupole isolation (1.6 m/z) and measurement in the Orbitrap (resolution 15,000 FWHM, fixed first mass 120 m/z). Fragmentation was performed after accumulation of 2x10^5^ ions or after filling time of 25 ms for each precursor ion (whichever occurred first). Only multiple charged (2+ −7+) precursor ions were selected for MS/MS. Dynamic exclusion was employed with maximum retention period of 30 s. Isotopes were excluded.

For data independent acquisition (DIA), 1 μg of reconstituted peptides were separated using a nanoAcquity UPLC (Waters, Milford, MA), which was coupled online to the MS. Peptide mixtures were separated in trap/elute mode, using a trapping (nanoAcquity Symmetry C18, 5 μm, 180 μm × 20 mm) and an analytical column (nanoAcquity BEH C18, 1.7 μm, 75 μm x 250 mm). The outlet of the latter was coupled directly to an Orbitrap Fusion Lumos mass spectrometer (Thermo Fisher Scientific, San Jose, CA) using the Proxeon nanospray source. Solvent A consisted of water, 0.1% formic acid and solvent B consisted of acetonitrile, 0.1% formic acid. The samples were loaded with a constant flow of solvent A, at 5 μL/min onto the trapping column with a trapping time of 6 min. Peptides were eluted via the analytical column at a constant flow of 300 nL/min. During this step, the percentage of solvent B increased in a nonlinear fashion from 0% to 40% within 120 min. Total runtime was 145 min, including cleanup and column re-equilibration. The peptides were introduced into the mass spectrometer via a Pico- Tip Emitter 360 µm OD x 20 µm ID; 10 µm tip (New Objective) and a spray voltage of 2.2 kV applied. The capillary temperature was set at 300 °C and the RF lens set to 30%. Full scan MS spectra with mass range 350-1650 m/z were acquired in profile mode in the Orbitrap at a resolution of 120,000 FWHM. The filling time was set at maximum of 20 ms with an AGC target of 5 x 10^5^ ions. DIA scans were acquired with 40 mass window segments of differing widths across the MS1 mass range and the HCD collision energy set to 30%. MS/MS scan resolution in the Orbitrap was set to 30,000 FWHM with a fixed first mass of 200m/z after accumulation of 1x 106 ions, or after filling time of 70 ms (whichever occurred first). Data were acquired in profile mode. Tune version 2.1 and Xcalibur 4.1 were used for data acquisition and processing. The MS/MS scan resolution in the Orbitrap was set to 30k with an AGC target of 1 × 106 and max injection time of 70 ms. All data acquisition was performed using XCalibur 4.0/Tune 2.1 (Thermo Fisher Scientific).

#### Mass spectrometry data analysis for DDA DIA samples

For DDA, data were analysed using MaxQuant (version 1.5.3.28) (Cox & Mann, 2008). MS/MS spectra were searched against the Mouse Swiss-Prot entries of the Uniprot KB (database release 2016_01, 16,756 entries), using the Andromeda search engine (Cox *et al*, 2011). A list of common contaminants was appended to the database search and the search criteria set as follows: full tryptic specificity was required (cleavage after lysine or arginine residues, unless followed by proline); 2 missed cleavages were allowed; oxidation (M) and acetylation (protein N-term) were applied as variable modifications, with mass tolerances of 20 ppm set for precursor and 0.5 Da for fragments. The reversed sequences of the target database were used as decoy database. Peptide and protein hits were filtered at a false discovery rate of 1% using a target-decoy strategy (Elias & Gygi, 2007). The protein group intensities per protein (from the proteinGroups.txt output of MaxQuant) were used for further analysis and visualization, both were performed with R (Team, 2019).

For DIA, acquired data were processed using Spectronaut Professional v13.10 (Biognosys AG). Raw files were searched using directDIA with Pulsar (Biognosys AG) against the mouse UniProt database (Mus musculus, entry only, release 2016_01), with a list of common contaminants appended, using default settings. Default Biognosys (BGS) factory settings were used for library generation. DIA data were searched using BGS factory settings, except: Proteotypicity Filter = Only Protein Group Specific; Major Group Quantity = Median peptide quantity; Major Group Top N = OFF; Minor Group Quantity = Median precursor quantity; Minor Group Top N = OFF; Data Filtering = Qvalue sparse; Normalization Strategy = Local normalization; Row Selection = Automatic. The candidates and protein report tables were exported from Spectronaut and used for further analysis as described below.

### Exploratory/Descriptive data analysis

All data analysis steps were completed with R (Team, 2019), unless otherwise stated. A PCA analysis and analysis of sample correlation was performed with the pcromp and cor function, respectively, to visualize the high dimensional data. The relevant comparisons were divided and specific abbreviations were assigned to each: A1 Wildtype stroke with NaCl treatment / Wildtype control; A2 Wildtype stroke with LPS priming / Wildtype control. Proteins were considered differentially expressed (“DEP”) when they were up- or down-regulated by a factor of 1.5 to the respective control value. This symmetrical definition of DEPs was ensured by transforming the fold changes to logarithmic values with a base of 2 (Quackenbush, 2002), resulting in a cut-off of │log2 FC│≥ 0.58. Furthermore, proteins with a q-value of ≤ 0.05 were considered as DEPs. Heatmaps were generated with the ComplexHeatmap package (Gu *et al*, 2016) in order to visualize different patterns and shared regulation of gene clusters. The protein quantities were z-score normalized for all samples by protein row-wise. The dendrogram was reordered with the R package dendsort (Sakai *et al*, 2014) for appropriate visualization. Correlation analysis of all and significantly changed proteins was achieved using the cor.test function from the R stats package and visualized by a customized function using the ggplot2 package. The correlation analysis was performed with the Spearman correlation method, since the Log2 fold changes of both datasets were not normally distributed. The intersections, as well as the uniquely changed DEPs, were retrieved by joining the A1 and A2 DEP lists with different functions from the dplyr package within the tidyverse package. These partitions were visualized with Venn diagrams using the draw.pairwise.venn function of the VennDiagram package (Chen & Boutros, 2011), before being subjected to a gene ontology and KEGG pathway enrichment analysis.

### Proteomic assessment of microglia enrichment

The proteomic data harvested from the CD11b+ and the mixed cell suspension containing all other brain cells (NTCF), was compared to a high-resolution proteomics dataset of oligodendrocytes, astrocytes, cortical neurons and microglia (Sharma *et al*., 2015). The cell-type specific proteome dataset was sorted for every protein to obtain the cell types showing the highest expression of the respective protein. Subsequently, the cell types showing the second highest expression were identified and a ratio for the respective protein between the expression values for these two cell types calculated. A protein was defined to be a microglia specific marker if this ratio showed at least a 5-fold higher expression in microglia than in the cell type with the second highest expression. This criterion was also applied to identify oligodendrocyte, astrocyte and cortical neuron specific marker proteins. The thus identified proteins of all other cell types were bound together for heatmap generation as “non-microglia proteins”. The obtained protein lists of cell-type specific marker proteins were then joined with the proteome of the CD11b+ samples and the mixed cell suspension to identify and count the number of cell-type specific proteins in the two different MACS fractions.

### Functional protein annotation and enrichment analyses

A gene enrichment and functional annotation analysis was performed by the DAVID platform (Huang da *et al*., 2009) with a conversion success rate of >99,9%. The gene lists, obtained by DEP analysis as well as Venn analysis, were submitted to DAVID and are illustrated in Table 1. The corresponding datasets of all proteins from the A1 and A2 analyses – or the intersections of all in both comparisons – detected proteins were used instead of the total *mus muscuclus* genome as background dataset for gene ontology enrichment analysis, in order to obtain specifically the enriched gene ontology terms within the determined microglia proteome. GO-terms that describe biological processes, GO-terms that describe cellular components and KEGG pathways were integrated into further analysis. The analysis was restricted to level 5 gene ontology (GO) terms representing a trade-off between high coverage of differentially expressed genes per GO-term and likewise a specific description of a biological process. A default threshold of at least 2 counts per GO-term was used. Lists of Gene Ontology terms of the biological processes with a Fisher’s Exact p-value ≤ 0.1 were either subjected to REVIGO analysis (Supek *et al*, 2011) for semantic similarity analysis and reduction, or analyzed using the simplifyEnrichment R package as described by the developers (Gu & Hubschmann, 2022). REVIGO analysis was achieved under the following parameters: medium list size (0.7) and mus musculus UniProt database as identifier with the by default preselected simRel score (Schlicker *et al*, 2006) used as semantic similarity measure. The resulting R scripts were modified to suit the way in which the graphical parameters should be displayed. Lists of enriched KEGG-pathways were filtered for a Fisher’s Exact p-value ≤ 0.05 and imported into R. A bubble plot was generated for visualization with a custom function using the ggplot2 package. Interesting enriched KEGG pathways were further visualized with the R package Pathview (Luo & Brouwer, 2013): Datasets were filtered for a q-value of ≤ 0.05 and relevant columns (Log2 Fold changes) of A1 and A2 were subsetted. The KEGG pathview figure of the respective pathway was downloaded and the proteomics data annotated via the gene symbols. The gene list of the gene ontology term “GO:0045335 phagocytic vesicle” was retrieved using the R package “org.Mm.eg.db” with a taxon filter for mus musculus.

**Table 1:** Protein lists for DAVID enrichment analysis.

|  | Up in A1 | Up in A2 | Down in A1 | Down in A2 |
| --- | --- | --- | --- | --- |
| A1_up | yes |  |  |  |
| A2_up |  | yes |  |  |
| A1_down |  |  | yes |  |
| A2_down |  |  |  | yes |
| A1A2upall | yes | yes | no | no |
| A1A2downall | no | no | yes | yes |
| A1_up_unique | yes | no | no |  |
| A2_up_unique | no | yes |  | no |
| A1_down_unique | no |  | yes | no |
| A2_down_unique |  | no | no | yes |

“yes” means that a protein from lists shown in the rows is also present in the list of the respective column; “no” means this is not the case and an empty box denotes no information can be stated as to whether or not a protein is present in the list of the respective column.

### Integration of other datasets for bioinformatic analyses

A dataset to determine the functional activation of microglia towards a pro- or anti-inflammatory state was retrieved, in which proteomic signatures of proinflammatory “classically activated” microglia and anti-inflammatory/“alternatively activated” microglia were determined (Bell-Temin *et al*., 2015). These gene lists were annotated to the list of proteins that were significantly regulated in either the A1 or A2 comparison via the UniProtIDs, to extract pro- and anti-inflammatory DEPs. The protein intensities of these proteins were z-score normalized, visualized with a heatmap and an analysis of differential protein abundance between protein sets belonging to the respective gene sets was conducted. Furthermore, a dataset of transcriptome profiles of isolated microglia exposed in vivo to Interferon-β was retrieved (Supplementary Dataset 3) (Deczkowska *et al*., 2017) and processed as described by the authors of the respective paper. The resultant list of 255 induced/upregulated genes in microglia by Interferon-β was provided a gene set for estimating the induction of a type I IFN responsive microglial state after tMCAO. Gene set enrichment analysis was performed using the fgsea R package (Korotkevich *et al*, 2021) and visualization p using the gggsea package (Huber). Bar plots and statistical testing were completed as described above and in the respective figure legends. The final task involved performing a cluster specific analysis of the type I IFN network as determined previously (Mostafavi *et al*., 2016). This dataset was obtained from the supplementary data (Table S3) to extract cluster wise the assigned regulator and target proteins. These gene sets were matched to all proteins from the A1 or A2 comparison. Subsequently, the joined datasets were filtered for a q-value of < 0.05 for network visualization. Differential protein abundances of cluster 3 regulator and target proteins between the different experimental groups were estimated via statistical verification on the paired datasets.

### Statistics

Data were presented as scatter dot plots, mean ± SEM. Shapiro-Wilk normality test was conducted before further statistical analysis was performed. Parametric (two-way ANOVA and repeated measures ANOVA with Post-hoc Holm-Sidak test, unpaired Student’s t-test with Welch’s correction) or non-parametric tests (Friedman-test with post-hoc Wilcoxon signed-rank test and adjustment of p-values for multiple testing with the Bonferroni-Holm correction) were completed where appropriate. Statistical tests for proteomic analysis and functional annotation and enrichment analyses were performed as indicated above. Differences were considered significant when p < 0.05. The statistical analysis was achieved using Graphpad Prism 8.4 or R (Team, 2019).

## Data availability

The mass spectrometry proteomics data have been deposited to the ProteomeXchange Consortium via the PRIDE partner repository (Perez-Riverol *et al*, 2022) with the dataset identifiers PXD031973 (Validation of microglia isolation protocol experiment) and PXD031930 (tMCAO experiment)

## Acknowledgments

The authors acknowledge Mrs. Rose-Marie Zimmer and Mr. Alexander Gloria for skillful technical assistance, Karol Szafranski and Philipp Koch for helpful bioinformatic and statistical advice and Leonie Karoline Stabenow for critical reading and editing of the manuscript. Open access funding provided by Projekt DEAL.

## Author contributions

Conceptualization: D-LH, RB; Data curation: D-LH, FH, EC, RB; Formal analysis: D-LH, FH, NR, TTDD, EKS, NO, LB; Investigation: D-LH, FH, RB; Methodology: D-LH, RB; Project administration: RB; Resources: RB, HM; Data analysis: D-LH, FH, EC, LB; Supervision: RB; Visualization: D-LH; Writing—original draft: D-LH, RB; Writing—review and editing: D-LH, HM, RB.

## Expanded View Figure legends

**Figure EV1:**
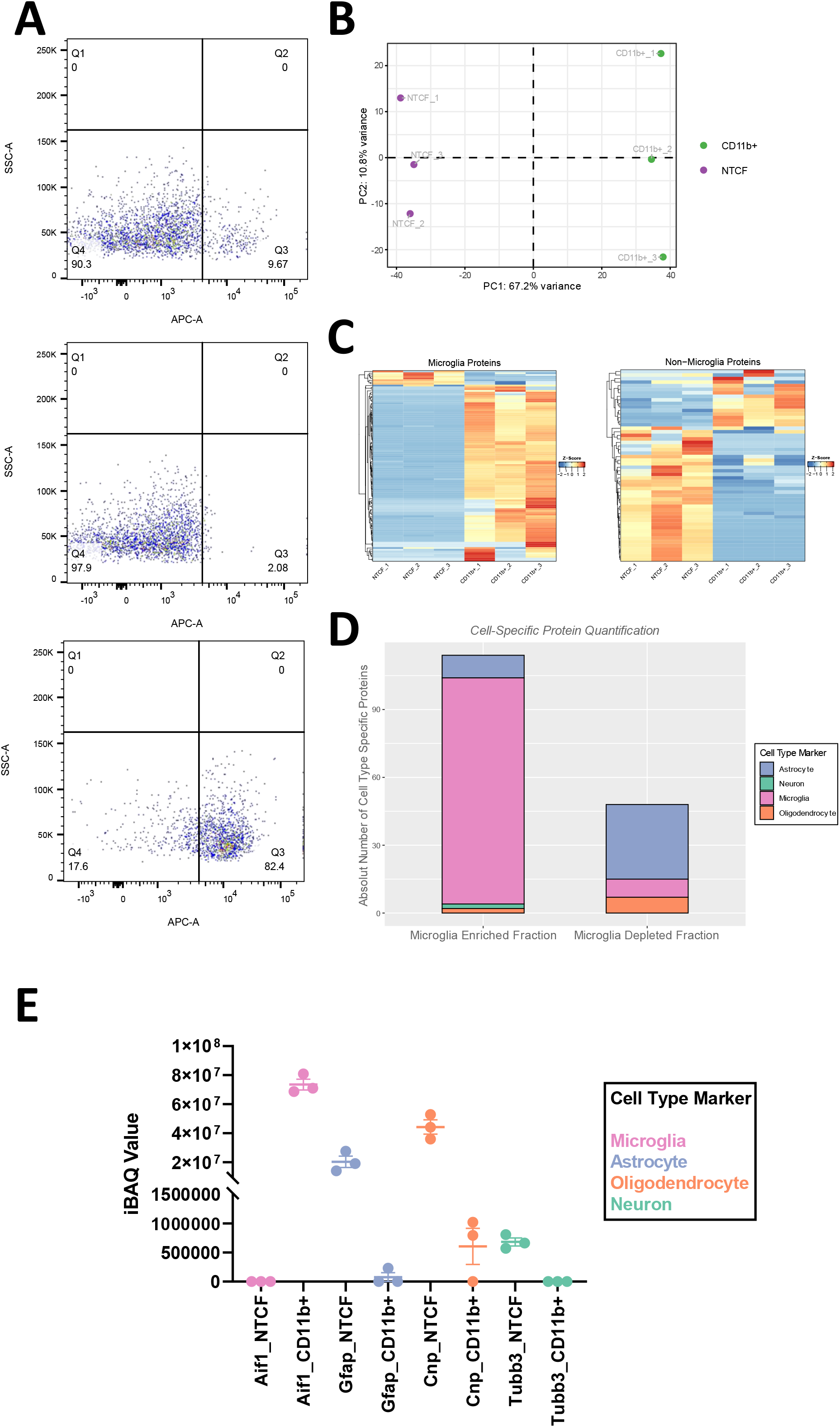
Validation of microglial cell enrichment via MACS. **A** FACS analysis of MACS-isolated CD11b+ microglia. Different cell suspensions before and after MACS-based separation were stained with an APC-coupled CD11b antibody, followed by FACS analysis. Upper plot: Representative FACS profile of the original fraction that still contains all brain cells before MACS separation. Middle plot: Representative FACS profile of the non-target cell fraction (NTCF) containing all CD11b- cells after MACS separation. Lower plot: Representative FACS profile of the CD11b positive cell fraction (CD11b+) containing all CD11b+ cells, mainly microglia. **B** Principal component analysis of proteomes determined for CD11b+ and NTCF fractions showing a clear separation of two clusters. **C** Proteins of cells from CD11b+ and NTCF fractions detected and quantified by proteomic analysis, were matched to a brain cell type proteome dataset by (Sharma *et al*., 2015). Heatmap visualization of all proteins defined to be marker proteins for microglial cells (left plot) and non-microglia cells (right plot). The z-score normalized protein intensities are depicted. A clear pattern of stronger protein intensities of non-microglia proteins within the NTCF samples and microglia proteins within the CD11b+ samples is visible (for both figures n=3 biological replicates sampled from the same mice, one CD11b+/NTCF pair per mouse). **D** Quantification of marker proteins for astrocytes, neurons, microglia, and oligodendrocytes found in the microglia-enriched fraction (CD11b+), or the microglia-depleted fraction (NTCF). Microglia marker proteins appear to be strongly enriched within the microglia-enriched fraction, while in the microglia-depleted fraction predominantly astrocyte and oligodendrocyte proteins were detected. **E** Dot plot chart of the log_10_(PG.Intensities) of classical neuronal (Tuj1) and glia marker proteins (Microglia: Iba1/Aif1, Astrocytes: Gfap, Oligodendrocytes: Cnp) in the proteome of CD11b+ and NTCF fractions. Iba1 was detected at high levels in CD11b+ samples only, while for all other marker proteins the opposite applied, i.e. high levels of other brain cell marker proteins were detected only in the NTCF fraction.

**Figure EV2:**
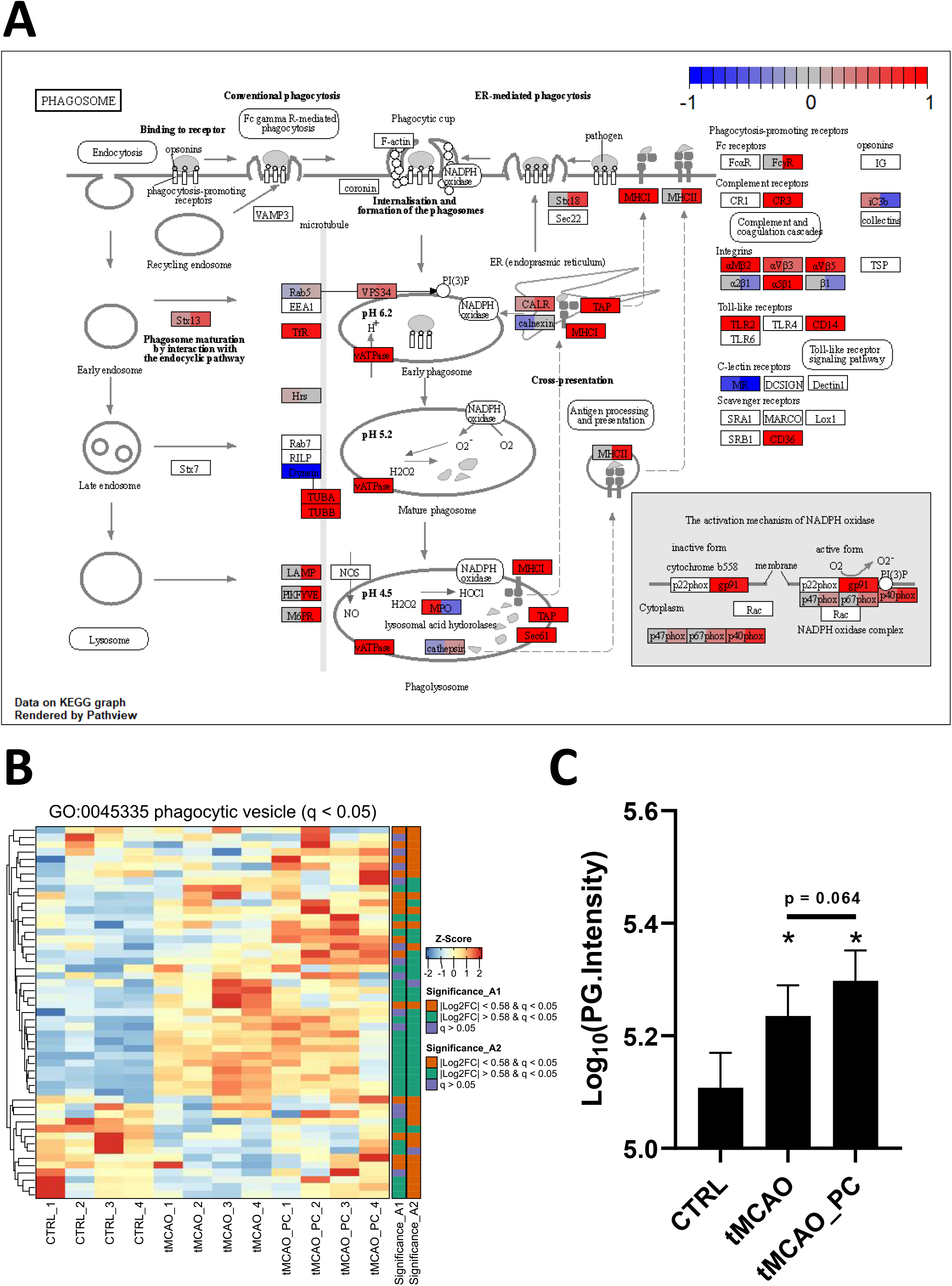
Induction of phagosome proteins and phagocytic vesicle-associated proteins after tMCAO or tMCAO and preconditioning. **A** Functional pathway annotation of significantly changed proteins in A1 and A2 to the KEGG pathway: Phagosome with the R package pathview (Luo & Brouwer, 2013). Log2FC of the re-spective proteins are normalized and shown on a scale between −1 and 1, where positive values indicating significant upregulation are colored red, downregulated proteins are colored blue. On the left part of the respective box with the respective protein name the normalized Log2FC of A1 is displayed; the right half of the respective box with the respective protein name displays the normalized Log2FC of A2. **B** Heatmap visualization of the z-score normalized protein intensities of all samples. Only proteins are shown that were significantly changed in A1 or A2 and belong to the GO-term “Phagocytic vesicle” (all samples n=4 biological replicates sampled from different mice). On the right side of the heatmap an annotation is shown as to whether a protein was significantly changed in A1 or A2 and if yes, above or below the fold change cut-off of │log2 FC│≥ 0.58. **C** Bar charts of Log10(PG.Intensities) of all in A1 or A2 significantly changed proteins belonging to the GO-term “Phagocytic vesicle” (n=51 proteins, a Repeated measures ANOVA was conducted, with a p-value < 0.0001 with Post-hoc Holm-Sidak tests with * = p < 0.05 for comparisons to the control group; specifically p = 0.0005 for CTRL vs. tMCAO, p < 0.0001 for CTRL vs. tMCAO_PC and p = 0.0640 for the comparison tMCAO to tMCAO_PC).

**Figure EV3:**
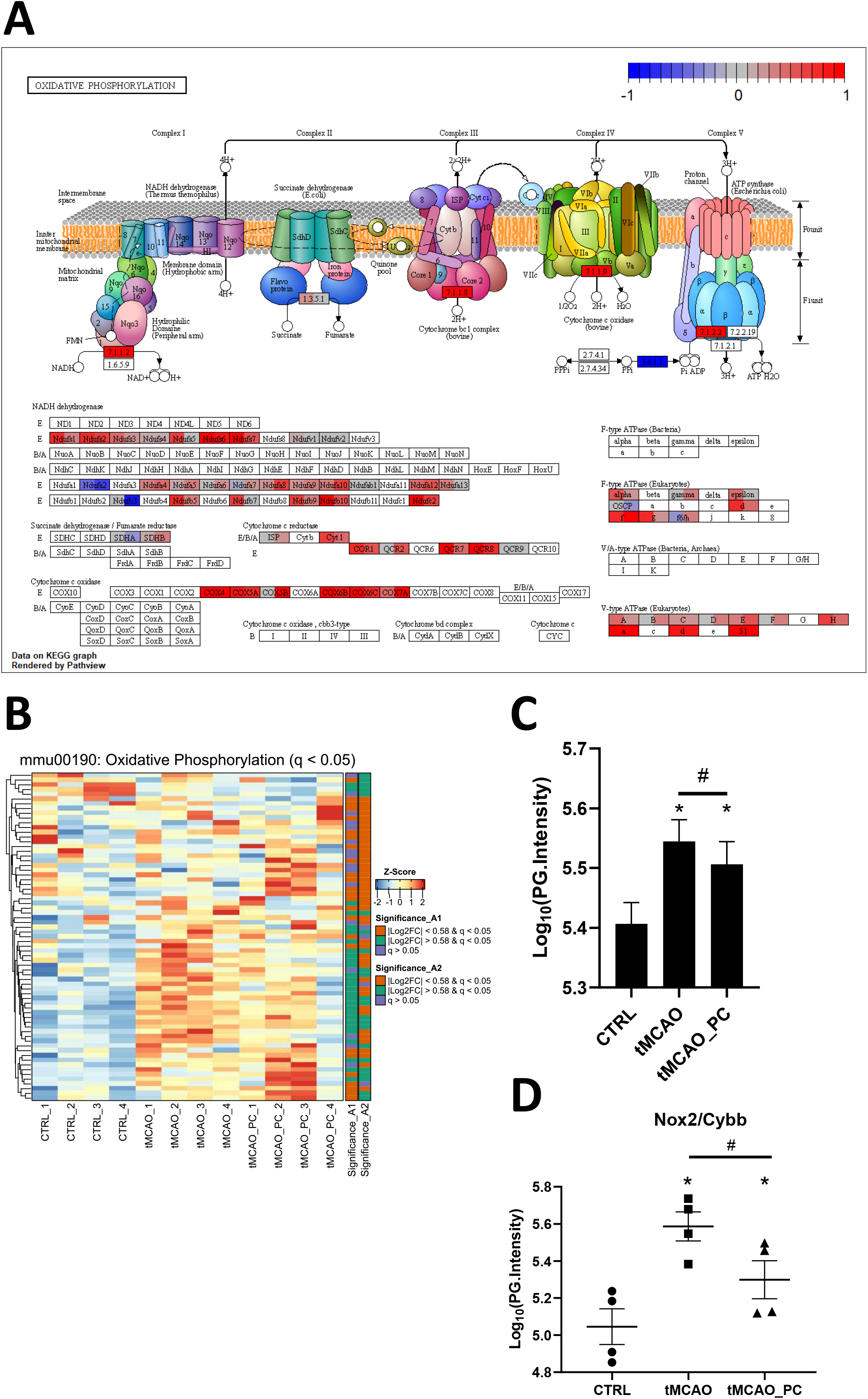
Amelioration of ROS-production pathways by preconditioning after tMCAO. **A** Functional pathway annotation of significantly changed proteins in A1 and A2 to the KEGG pathway mmu00190: “Oxidative Phosphorylation” with the R package pathview (Luo & Brouwer, 2013). Log2FC of the respective proteins are normalized and shown on a scale between −1 and 1, where positive values indicating significant upregulation are colored red, downregulated proteins are colored blue. On the left half of the respective box with the respective protein name, the normalized Log2FC of A1 is displayed; the right half of the respective box with the respective protein name displays the normalized Log2FC of A2. **B** Heatmap visualization of the z-score normalized protein intensities of all samples. Only proteins are shown that were significantly changed in A1 or A2 and belong to the KEGG pathway mmu00190: “Oxidative Phosphorylation” (all samples n=4 biological replicates sampled from different mice). On the right side of the heatmap an annotation is shown as to whether a protein was significantly changed in A1 or A2 and if yes, above or below the fold change cut-off of │log2 FC│≥ 0.58. **C** Bar charts of Log10(PG.Intensities) of all significantly changed proteins in A1 or A2 belonging to the KEGG pathway mmu00190: “Oxidative Phosphorylation” (n=69 proteins, a Repeated measures ANOVA was conducted, with a p-value < 00001 with Post-hoc Holm-Sidak tests with * = p < 0.05 for comparisons to the control group; specifically p < 0.0001 for both comparisons CTRL vs. tMCAO and CTRL vs. tMCAO_PC and # = p < 0.05 for the comparison tMCAO to tMCAO_PC, specifically p = 0.0424). **D** Bar chart of Log10(PG.Intensities) of Nox2 (n=4, each). Shown are the FDR-adjusted p-values (=q-values) as reported by Spectronaut software in the differential abundance analysis; *q ≤ 0.05 for comparisons to the control group and ^#^ q ≤ 0.05 for the comparison tMCAO to tMCAO_PC. Specifically, q-values were q = 0.000929 for tMCAO vs. CTRL, q = 0.000903 for tMCAO_PC vs. CTRL and q = 0.0044 for tMCAO_PC vs. tMCAO

**Figure EV4:**
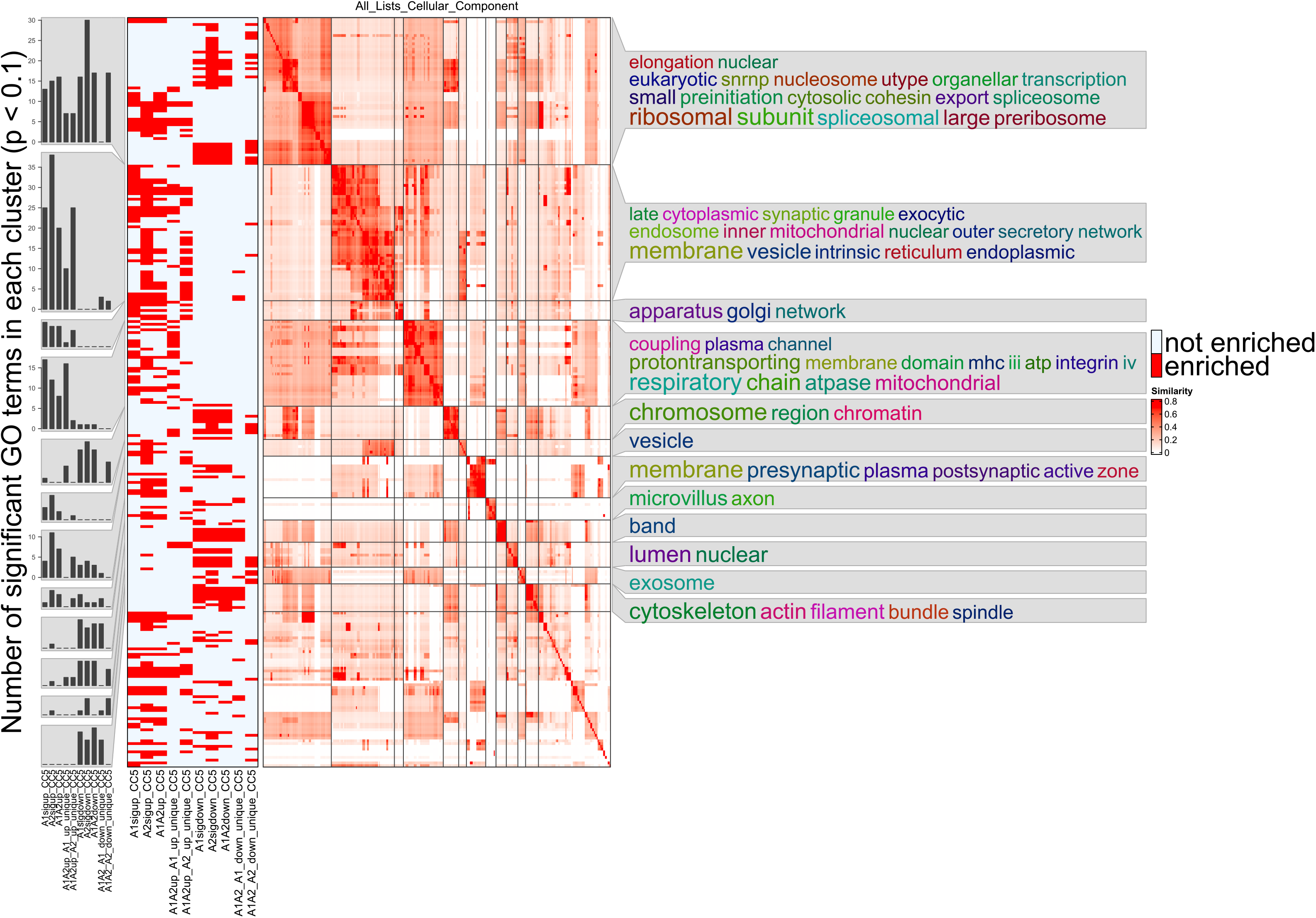
Gene ontology enrichment analysis for cellular components. Functional gene ontology enrichment analysis of level 5 GO-terms of cellular components was conducted based on the experimental groups A1/A2_up/down, standing for the up- and downregulated groups from the single comparisons tMCAO vs. CTRL and tMCAO_PC vs. CTRL, and thereafter the protein lists derived from the Venn diagrams. Enriched GO-terms with a Fisher’s Exact p-value ≤ 0.1 were used for semantic similarity clustering – the similarity matrix with specific clusters is depicted in the middle. Each row corresponds to one GO-term and in the matrix each column corresponds to one GO-term. On the left a binary heatmap is displayed showing whether a GO-term was enriched in the protein target list or not, where each row corresponds to one GO-term. On the extreme left, bar plots are shown where clusterwise the number of GO-terms for each protein list is counted. On the right a word cloud is annotated to the similarity matrix, providing information about the cellular component in keywords, in terms of an overrepresentation of individual words from all GO-terms located in this cluster.

**Figure EV5:**
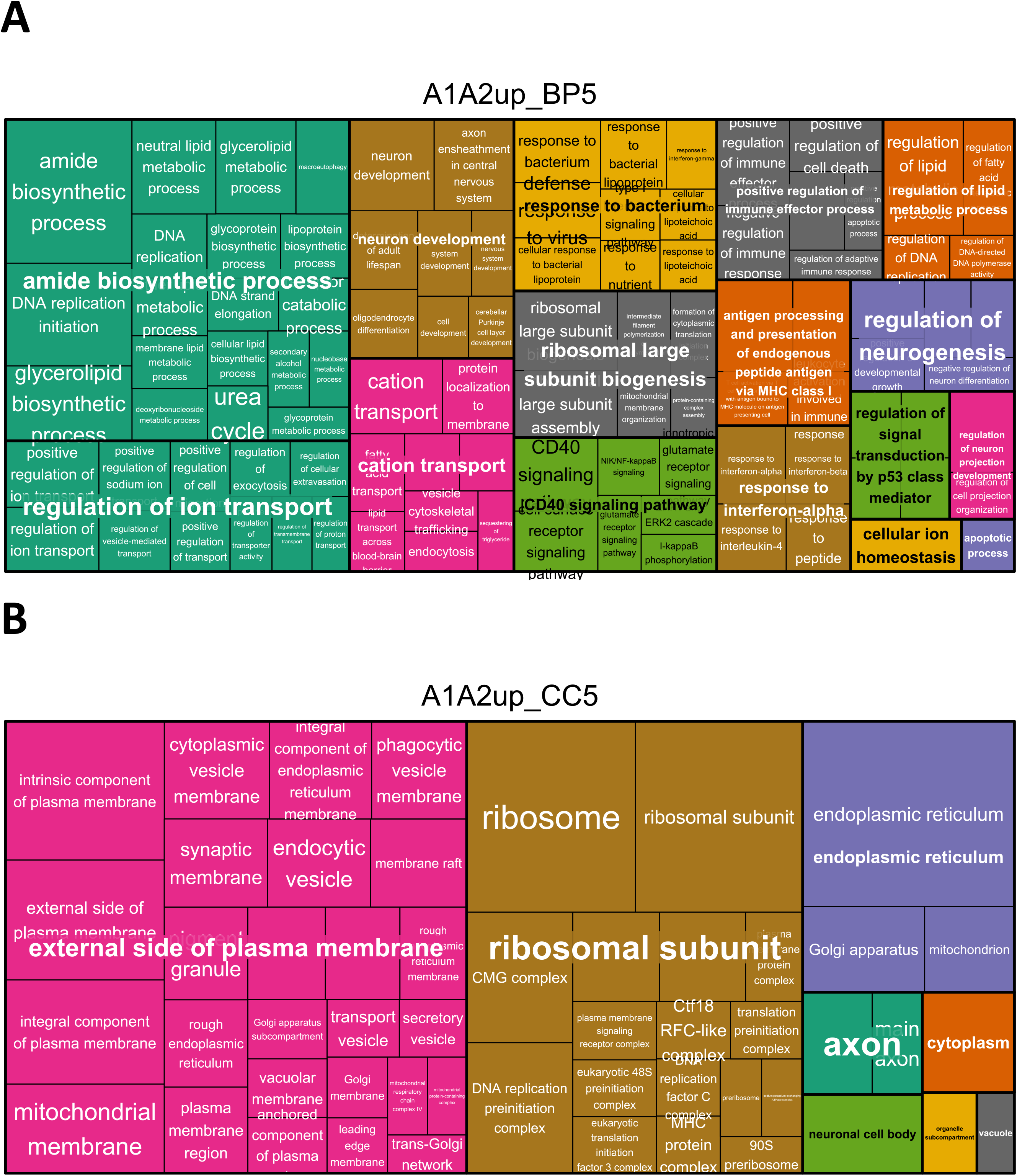
Treemaps of biological process GO-terms enriched in with tMCAO and tMCAO_PC upregulated microglia proteins. **A** Level 5 GO-terms of biological processes enriched in the lists of proteins upregulated in both comparisons (A1&A2) are visualized within a treemap generated with Revigo. Semantically different clusters are colored differentially. The cluster size is a function of the Fisher’s Exact p-values of the respective enriched GO-clusters and GO-terms: if the p-value is smaller, the cluster size is bigger. All GO-terms were filtered for a cut-off of Fisher’s Exact p-value ≤ 0.1. **B** Level 5 GO-terms of cellular components enriched in the lists of proteins upregulated in both comparisons (A1&A2) are visualized within a treemap generated with Revigo. Semantically similar clusters are colored differentially. The cluster size is a function of the Fisher’s Exact p- values of the respective enriched GO-clusters and GO-terms: if the p-value is smaller, the cluster size is bigger. All GO-terms were filtered for a cut-off of Fisher’s Exact p-value ≤ 0.1.

**Figure EV6:**
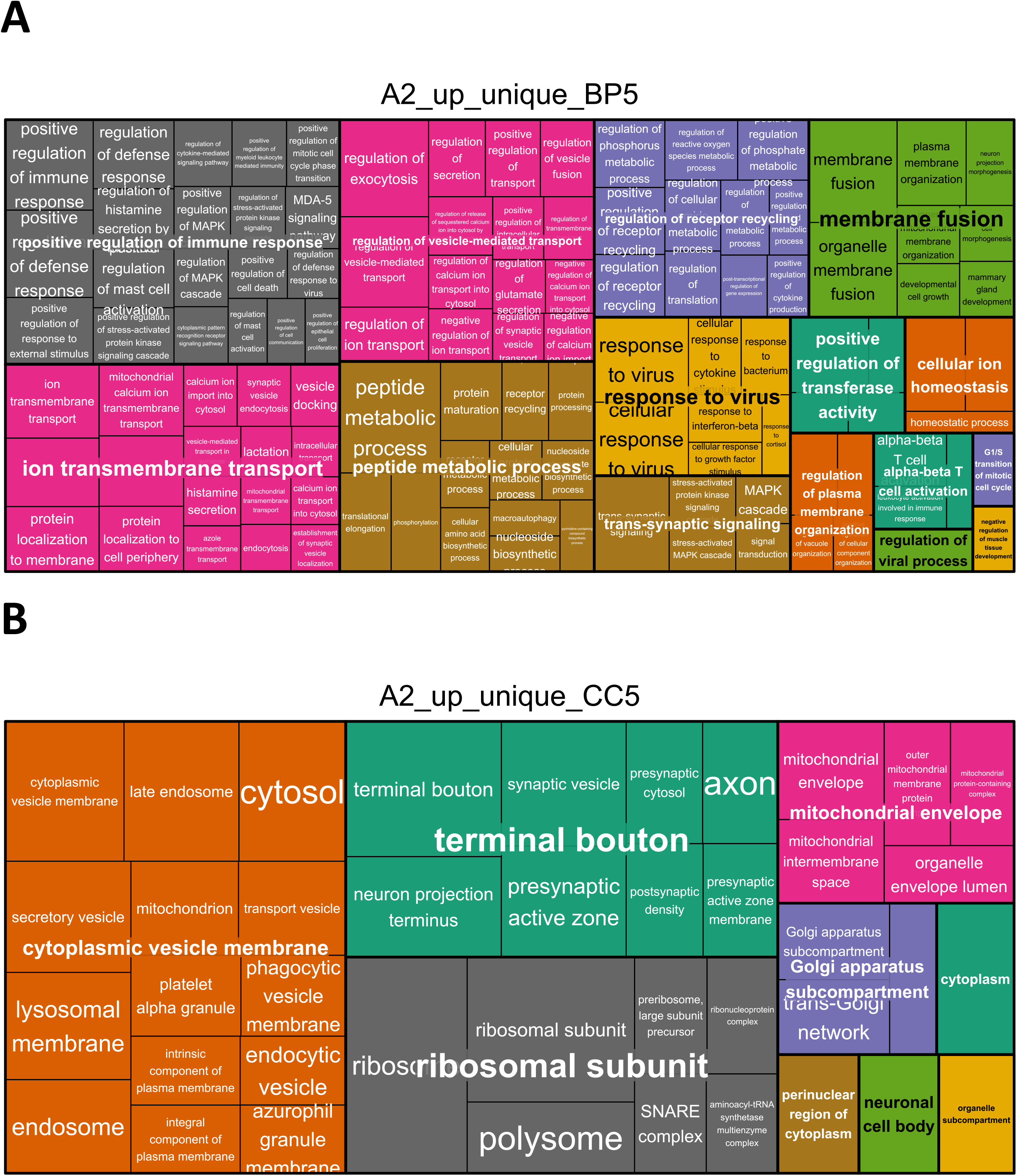
Treemaps of biological process GO-terms enriched uniquely in with tMCAO_PC upregulated microglia proteins. **A** Level 5 GO-terms of biological processes enriched in the lists of proteins upregulated uniquely in LPS-primed microglia after tMCAO (A2_up_unique) are visualized within a treemap generated with Revigo. Semantically similar clusters are colored differentially. The cluster size is a function of the Fisher’s Exact p-values of the respective enriched GO-clusters and GO-terms: if the p-value is smaller, the cluster size is bigger. All GO-terms were filtered for a cut-off of Fisher’s Exact p-value ≤ 0.1. **B** Level 5 GO-terms of cellular components enriched in the lists of proteins upregulated uniquely in LPS-primed microglia after tMCAO (A2_up_unique) are visualized within a treemap generated with Revigo. Semantically similar clusters are colored differentially. The cluster size is a function of the Fisher’s Exact p-values of the respective enriched GO-clusters and GO-terms: if the p-value is smaller, the cluster size is bigger. All GO-terms were filtered for a cut-off of Fisher’s Exact p-value ≤ 0.1.

**Figure EV7:**
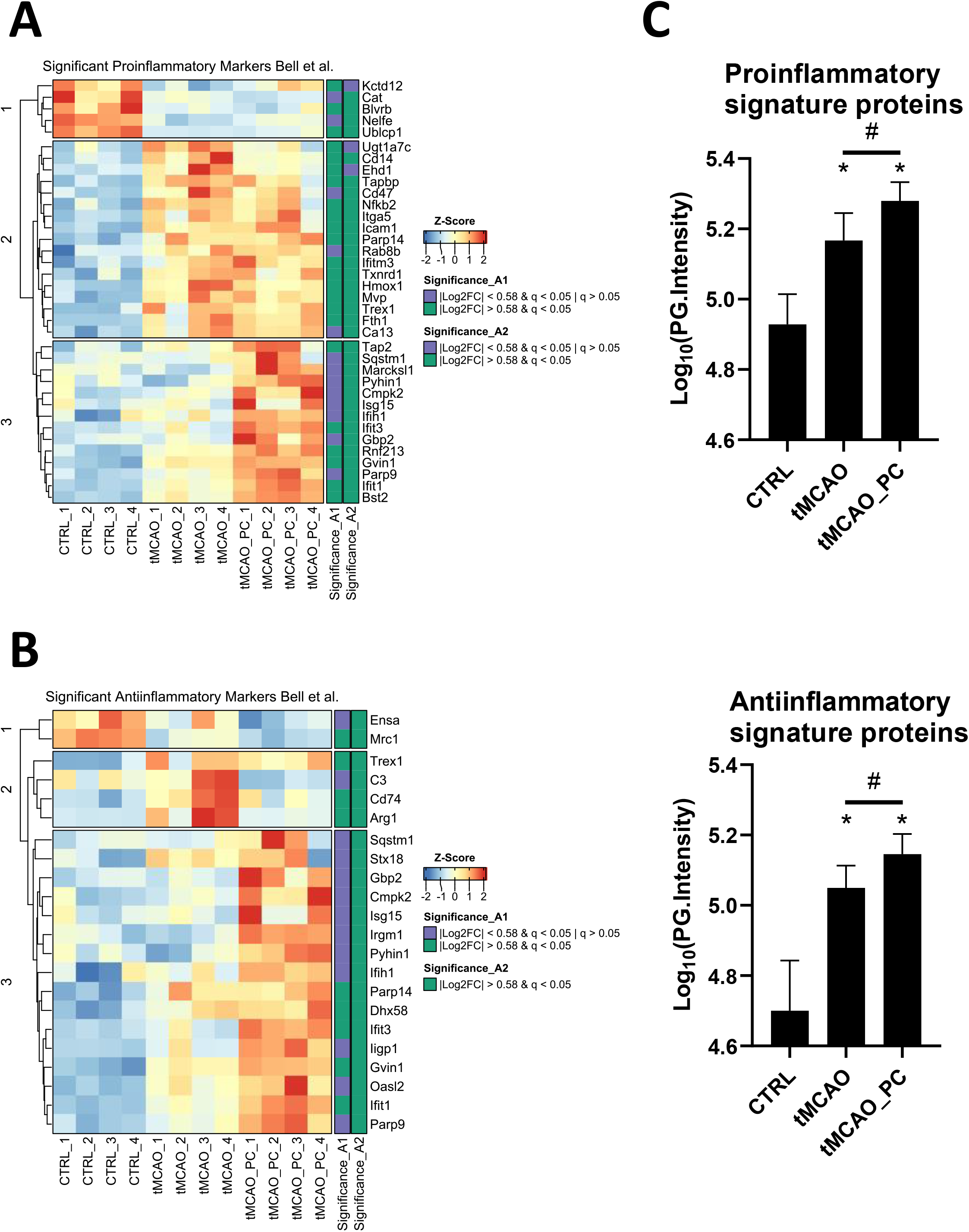
Proteomic signature of enhanced inflammatory activation of microglia after tMCAO and preceding LPS-induced inflammatory preconditioning. **A, B** Heatmap visualizations of the z-score normalized protein intensities of all samples. Proteins are shown when changed in A1 or A2 (│log2 FC│≥ 0.58 and q-value ≤ 0.05) and belong to the corresponding microglial pro-/anti-inflammatory signature (Bell-Temin *et al*., 2015) (n=4, each). On the right side of the heatmap an annotation is shown as to whether a protein was significantly changed in A1 or A2 above or below the fold change cut-off of │log2 FC│≥ 0.58. **C** Bar charts of Log10 (PG.Intensities, where PG = protein group) of all in A1 or A2 DEPs belonging to either the pro- (upper chart) or anti-inflammatory (lower chart) gene signatures (n=36 proteins for proinflammatory gene signature, n=22 proteins for anti-inflammatory gene signature. A Friedman-test was conducted, with a p-value < 0.05 for both signatures, specifically p < 0.0001 for the proinflammatory signature and p = 0.0002 for the anti-inflammatory signature. Post-hoc Wilcoxon signed-rank tests were conducted with *p < 0.05 for comparisons to the control group and ^#^p < 0.05 for the comparison tMCAO to tMCAO_PC. Specifically, p-values for the proinflammatory signature were p = 0.0003 for both CTRL vs. tMCAO and CTRL vs. tMCAO_PC and p = 0.0105 for tMCAO vs. tMCAO_PC. For the anti-inflammatory signature p-values were p = 0.0014 for CTRL vs. tMCAO, p = 0.003 for CTRL vs. tMCAO_PC and p = 0.0462 for tMCAO vs. tMCAO_PC.) All p-values were adjusted for multiple testing with Bonferroni-Holm correction.

## SUPPLEMENTARY TABLES

### SUPPLEMENTARY TABLE EV1

**Table EV 1:**
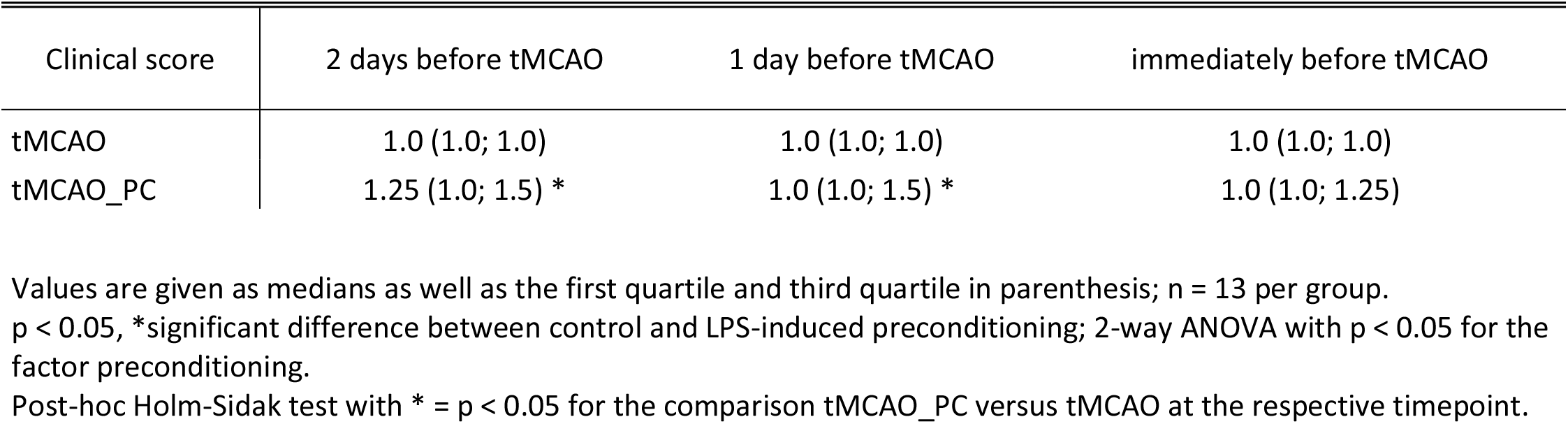
Effect of LPS preconditioning (0.8 µg/g b.w., i.p.) on clinical course before tMCAO administration.

## References

Bederson JB, Pitts LH, Tsuji M, Nishimura MC, Davis RL, Bartkowski H (1986) Rat middle cerebral artery occlusion: evaluation of the model and development of a neurologic examination. Stroke 17: 472–476

Bell-Temin H, Culver-Cochran AE, Chaput D, Carlson CM, Kuehl M, Burkhardt BR, Bickford PC, Liu B, Stevens SM, Jr. (2015) Novel Molecular Insights into Classical and Alternative Activation States of Microglia as Revealed by Stable Isotope Labeling by Amino Acids in Cell Culture (SILAC)-based Proteomics. Mol Cell Proteomics 14: 3173–3184

Bennett ML, Bennett FC, Liddelow SA, Ajami B, Zamanian JL, Fernhoff NB, Mulinyawe SB, Bohlen CJ, Adil A, Tucker A et al (2016) New tools for studying microglia in the mouse and human CNS. Proc Natl Acad Sci U S A 113: E1738–1746

Bonati LH, Jongen LM, Haller S, Flach HZ, Dobson J, Nederkoorn PJ, Macdonald S, Gaines PA, Waaijer A, Stierli P et al (2010) New ischaemic brain lesions on MRI after stenting or endarterectomy for symptomatic carotid stenosis: a substudy of the International Carotid Stenting Study (ICSS). Lancet Neurol 9: 353–362

Bordet R, Deplanque D, Maboudou P, Puisieux F, Pu Q, Robin E, Martin A, Bastide M, Leys D, Lhermitte M et al (2000) Increase in endogenous brain superoxide dismutase as a potential mechanism of lipopolysaccharide-induced brain ischemic tolerance. J Cereb Blood Flow Metab 20: 1190–1196

Chamorro A, Lo EH, Renu A, van Leyen K, Lyden PD (2021) The future of neuroprotection in stroke. J Neurol Neurosurg Psychiatry 92: 129–135

Chen H, Boutros PC (2011) VennDiagram: a package for the generation of highly-customizable Venn and Euler diagrams in R. BMC Bioinformatics 12: 35

Collaborators GBDS (2019) Global, regional, and national burden of stroke, 1990-2016: a systematic analysis for the Global Burden of Disease Study 2016. Lancet Neurol 18: 439–458

Cox J, Mann M (2008) MaxQuant enables high peptide identification rates, individualized p.p.b.-range mass accuracies and proteome-wide protein quantification. Nat Biotechnol 26: 1367–1372

Cox J, Neuhauser N, Michalski A, Scheltema RA, Olsen JV, Mann M (2011) Andromeda: a peptide search engine integrated into the MaxQuant environment. J Proteome Res 10: 1794–1805

Davalos D, Grutzendler J, Yang G, Kim JV, Zuo Y, Jung S, Littman DR, Dustin ML, Gan WB (2005) ATP mediates rapid microglial response to local brain injury in vivo. Nat Neurosci 8: 752–758

Deczkowska A, Matcovitch-Natan O, Tsitsou-Kampeli A, Ben-Hamo S, Dvir-Szternfeld R, Spinrad A, Singer O, David E, Winter DR, Smith LK et al (2017) Mef2C restrains microglial inflammatory response and is lost in brain ageing in an IFN-I-dependent manner. Nat Commun 8: 717

DeGracia DJ (2017) Regulation of mRNA following brain ischemia and reperfusion. Wiley Interdiscip Rev RNA 8

Dirnagl U, Becker K, Meisel A (2009) Preconditioning and tolerance against cerebral ischaemia: from experimental strategies to clinical use. Lancet Neurol 8: 398–412

Dirnagl U, Simon RP, Hallenbeck JM (2003) Ischemic tolerance and endogenous neuroprotection. Trends Neurosci 26: 248–254

Ekker MS, Verhoeven JI, Vaartjes I, van Nieuwenhuizen KM, Klijn CJM, de Leeuw FE (2019) Stroke incidence in young adults according to age, subtype, sex, and time trends. Neurology 92: e2444–e2454

El Khoury J, Toft M, Hickman SE, Means TK, Terada K, Geula C, Luster AD (2007) Ccr2 deficiency impairs microglial accumulation and accelerates progression of Alzheimer-like disease. Nat Med 13: 432–438

Elias JE, Gygi SP (2007) Target-decoy search strategy for increased confidence in large-scale protein identifications by mass spectrometry. Nat Methods 4: 207–214

Garcia-Bonilla L, Benakis C, Moore J, Iadecola C, Anrather J (2014) Immune mechanisms in cerebral ischemic tolerance. Front Neurosci 8: 44

Gesuete R, Packard AE, Vartanian KB, Conrad VK, Stevens SL, Bahjat FR, Yang T, Stenzel-Poore MP (2012) Poly-ICLC preconditioning protects the blood-brain barrier against ischemic injury in vitro through type I interferon signaling. J Neurochem 123 Suppl 2: 75–85

Gidday JM (2006) Cerebral preconditioning and ischaemic tolerance. Nat Rev Neurosci 7: 437–448

Gordon CJ, Becker P, Ali JS (1998) Behavioral thermoregulatory responses of single- and group-housed mice. Physiol Behav 65: 255–262

Gu Z, Eils R, Schlesner M (2016) Complex heatmaps reveal patterns and correlations in multidimensional genomic data. Bioinformatics 32: 2847–2849

Gu Z, Hubschmann D (2022) Simplify enrichment: A bioconductor package for clustering and visualizing functional enrichment results. Genomics Proteomics Bioinformatics

Hamner MA, McDonough A, Gong DC, Todd LJ, Rojas G, Hodecker S, Ransom CB, Reh TA, Ransom BR, Weinstein JR (2022) Microglial depletion abolishes ischemic preconditioning in white matter. Glia 70: 661–674

Hamner MA, Ye Z, Lee RV, Colman JR, Le T, Gong DC, Ransom BR, Weinstein JR (2015) Ischemic Preconditioning in White Matter: Magnitude and Mechanism. J Neurosci 35: 15599–15611

Harris MA, Clark J, Ireland A, Lomax J, Ashburner M, Foulger R, Eilbeck K, Lewis S, Marshall B, Mungall C et al (2004) The Gene Ontology (GO) database and informatics resource. Nucleic Acids Res 32: D258–261

Hickman SE, Kingery ND, Ohsumi TK, Borowsky ML, Wang LC, Means TK, El Khoury J (2013) The microglial sensome revealed by direct RNA sequencing. Nat Neurosci 16: 1896–1905

Hochrainer K, Yang W (2022) Stroke Proteomics: From Discovery to Diagnostic and Therapeutic Applications. Circ Res 130: 1145–1166

Horgan RP, Kenny LC (2011) SAC review ‘Omic’ technologies: genomics, transcriptomics, proteomics and metabolomics. Obstet Gynaecol 13: 189–195

Hosmane S, Tegenge MA, Rajbhandari L, Uapinyoying P, Ganesh Kumar N, Thakor N, Venkatesan A (2012) Toll/interleukin-1 receptor domain-containing adapter inducing interferon-beta mediates microglial phagocytosis of degenerating axons. J Neurosci 32: 7745–7757

Huang da W, Sherman BT, Lempicki RA (2009) Systematic and integrative analysis of large gene lists using DAVID bioinformatics resources. Nat Protoc 4: 44–57

Huang Z, Huang PL, Panahian N, Dalkara T, Fishman MC, Moskowitz MA (1994) Effects of cerebral ischemia in mice deficient in neuronal nitric oxide synthase. Science 265: 1883–1885

Huber N, gggsea https://github.com/NicolasH2/gggsea.

Ivashkiv LB, Donlin LT (2014) Regulation of type I interferon responses. Nat Rev Immunol 14: 36–49

Jayaraj RL, Azimullah S, Beiram R, Jalal FY, Rosenberg GA (2019) Neuroinflammation: friend and foe for ischemic stroke. J Neuroinflammation 16: 142

Kariko K, Weissman D, Welsh FA (2004) Inhibition of toll-like receptor and cytokine signaling--a unifying theme in ischemic tolerance. J Cereb Blood Flow Metab 24: 1288–1304

Kirino T (2002) Ischemic tolerance. J Cereb Blood Flow Metab 22: 1283–1296

Korotkevich G, Sukhov V, Budin N, Shpak B, Artyomov MN, Sergushichev A (2021) Fast gene set enrichment analysis. bioRxiv: 060012

Kunz A, Park L, Abe T, Gallo EF, Anrather J, Zhou P, Iadecola C (2007) Neurovascular protection by ischemic tolerance: role of nitric oxide and reactive oxygen species. J Neurosci 27: 7083–7093

Kuo PC, Scofield BA, Yu IC, Chang FL, Ganea D, Yen JH (2016) Interferon-beta Modulates Inflammatory Response in Cerebral Ischemia. J Am Heart Assoc 5

Kurtin PJ, Bonin DM (1994) Immunohistochemical demonstration of the lysosome-associated glycoprotein CD68 (KP-1) in granular cell tumors and schwannomas. Hum Pathol 25: 1172–1178

Lajqi T, Köstlin-Gille N, Bauer R, Zarogiannis SG, Esra, Lajqi E, Ajeti V, 4 Stefanie Dietz SSA, Kranig SA, Rühle J, Demaj A, et al (2023 subm) Training vs. tolerance – the yin/yang of the innate immune system.

Lajqi T, Lang GP, Haas F, Williams DL, Hudalla H, Bauer M, Groth M, Wetzker R, Bauer R (2019) Memory-Like Inflammatory Responses of Microglia to Rising Doses of LPS: Key Role of PI3Kgamma. Front Immunol 10: 2492

Lajqi T, Marx C, Hudalla H, Haas F, Grosse S, Wang ZQ, Heller R, Bauer M, Wetzker R, Bauer R (2021) The Role of the Pathogen Dose and PI3Kgamma in Immunometabolic Reprogramming of Microglia for Innate Immune Memory. Int J Mol Sci 22

Leung PY, Stevens SL, Packard AE, Lessov NS, Yang T, Conrad VK, van den Dungen NN, Simon RP, Stenzel-Poore MP (2012) Toll-like receptor 7 preconditioning induces robust neuroprotection against stroke by a novel type I interferon-mediated mechanism. Stroke 43: 1383–1389

Lind L, Siegbahn A, Lindahl B, Stenemo M, Sundstrom J, Arnlov J (2015) Discovery of New Risk Markers for Ischemic Stroke Using a Novel Targeted Proteomics Chip. Stroke 46: 3340–3347

Luo W, Brouwer C (2013) Pathview: an R/Bioconductor package for pathway-based data integration and visualization. Bioinformatics 29: 1830–1831

Maier CM, Yu F, Nishi T, Lathrop SJ, Chan PH (2006) Interferon-beta fails to protect in a model of transient focal stroke. Stroke 37: 1116–1119

Marsh B, Stevens SL, Packard AE, Gopalan B, Hunter B, Leung PY, Harrington CA, Stenzel-Poore MP (2009) Systemic lipopolysaccharide protects the brain from ischemic injury by reprogramming the response of the brain to stroke: a critical role for IRF3. J Neurosci 29: 9839–9849

McDonough A, Lee RV, Noor S, Lee C, Le T, Iorga M, Phillips JLH, Murphy S, Moller T, Weinstein JR (2017) Ischemia/Reperfusion Induces Interferon-Stimulated Gene Expression in Microglia. J Neurosci 37: 8292–8308

McDonough A, Noor S, Lee RV, Dodge R, 3rd, Strosnider JS, Shen J, Davidson S, Moller T, Garden GA, Weinstein JR (2020) Ischemic preconditioning induces cortical microglial proliferation and a transcriptomic program of robust cell cycle activation. Glia 68: 76–94

McDonough A, Weinstein JR (2016) Neuroimmune Response in Ischemic Preconditioning. Neurotherapeutics 13: 748–761

McDonough A, Weinstein JR (2020) The role of microglia in ischemic preconditioning. Glia 68: 455–471

Mostafavi S, Yoshida H, Moodley D, LeBoite H, Rothamel K, Raj T, Ye CJ, Chevrier N, Zhang SY, Feng T et al (2016) Parsing the Interferon Transcriptional Network and Its Disease Associations. Cell 164: 564–578

Nair S, Sobotka KS, Joshi P, Gressens P, Fleiss B, Thornton C, Mallard C, Hagberg H (2019) Lipopolysaccharide-induced alteration of mitochondrial morphology induces a metabolic shift in microglia modulating the inflammatory response in vitro and in vivo. Glia 67: 1047–1061

Neumann H, Kotter MR, Franklin RJ (2009) Debris clearance by microglia: an essential link between degeneration and regeneration. Brain 132: 288–295

Nimmerjahn A, Kirchhoff F, Helmchen F (2005) Resting microglial cells are highly dynamic surveillants of brain parenchyma in vivo. Science 308: 1314–1318

Nogueira RG, Jadhav AP, Haussen DC, Bonafe A, Budzik RF, Bhuva P, Yavagal DR, Ribo M, Cognard C, Hanel RA et al (2018) Thrombectomy 6 to 24 Hours after Stroke with a Mismatch between Deficit and Infarct. N Engl J Med 378: 11–21

Obernier JA, Baldwin RL (2006) Establishing an appropriate period of acclimatization following transportation of laboratory animals. ILAR J 47: 364–369

Perez-Riverol Y, Bai J, Bandla C, Garcia-Seisdedos D, Hewapathirana S, Kamatchinathan S, Kundu DJ, Prakash A, Frericks-Zipper A, Eisenacher M et al (2022) The PRIDE database resources in 2022: a hub for mass spectrometry-based proteomics evidences. Nucleic Acids Res 50: D543–D552

Quackenbush J (2002) Microarray data normalization and transformation. Nat Genet 32 Suppl: 496–501

Rajbhandari L, Tegenge MA, Shrestha S, Ganesh Kumar N, Malik A, Mithal A, Hosmane S, Venkatesan A (2014) Toll-like receptor 4 deficiency impairs microglial phagocytosis of degenerating axons. Glia 62: 1982–1991

Rosenzweig HL, Lessov NS, Henshall DC, Minami M, Simon RP, Stenzel-Poore MP (2004) Endotoxin preconditioning prevents cellular inflammatory response during ischemic neuroprotection in mice. Stroke 35: 2576–2581

Rosenzweig HL, Minami M, Lessov NS, Coste SC, Stevens SL, Henshall DC, Meller R, Simon RP, Stenzel-Poore MP (2007) Endotoxin preconditioning protects against the cytotoxic effects of TNFalpha after stroke: a novel role for TNFalpha in LPS-ischemic tolerance. J Cereb Blood Flow Metab 27: 1663–1674

Sakai R, Winand R, Verbeiren T, Moere AV, Aerts J (2014) dendsort: modular leaf ordering methods for dendrogram representations in R. F1000Res 3: 177

Schilling M, Besselmann M, Muller M, Strecker JK, Ringelstein EB, Kiefer R (2005) Predominant phagocytic activity of resident microglia over hematogenous macrophages following transient focal cerebral ischemia: an investigation using green fluorescent protein transgenic bone marrow chimeric mice. Exp Neurol 196: 290–297

Schlicker A, Domingues FS, Rahnenfuhrer J, Lengauer T (2006) A new measure for functional similarity of gene products based on Gene Ontology. BMC Bioinformatics 7: 302

Schmidt C, Frahm C, Schneble N, Muller JP, Brodhun M, Franco I, Witte OW, Hirsch E, Wetzker R, Bauer R (2016) Phosphoinositide 3-Kinase gamma Restrains Neurotoxic Effects of Microglia After Focal Brain Ischemia. Mol Neurobiol 53: 5468–5479

Sharma K, Schmitt S, Bergner CG, Tyanova S, Kannaiyan N, Manrique-Hoyos N, Kongi K, Cantuti L, Hanisch UK, Philips MA et al (2015) Cell type- and brain region-resolved mouse brain proteome. Nat Neurosci 18: 1819–1831

Simpson DSA, Oliver PL (2020) ROS Generation in Microglia: Understanding Oxidative Stress and Inflammation in Neurodegenerative Disease. Antioxidants (Basel) 9

Stenzel-Poore MP, Stevens SL, King JS, Simon RP (2007) Preconditioning reprograms the response to ischemic injury and primes the emergence of unique endogenous neuroprotective phenotypes: a speculative synthesis. Stroke 38: 680–685

Stetler RA, Leak RK, Gan Y, Li P, Zhang F, Hu X, Jing Z, Chen J, Zigmond MJ, Gao Y (2014) Preconditioning provides neuroprotection in models of CNS disease: paradigms and clinical significance. Prog Neurobiol 114: 58–83

Stevens SL, Ciesielski TM, Marsh BJ, Yang T, Homen DS, Boule JL, Lessov NS, Simon RP, Stenzel-Poore MP (2008) Toll-like receptor 9: a new target of ischemic preconditioning in the brain. J Cereb Blood Flow Metab 28: 1040–1047

Stevens SL, Leung PY, Vartanian KB, Gopalan B, Yang T, Simon RP, Stenzel-Poore MP (2011) Multiple preconditioning paradigms converge on interferon regulatory factor-dependent signaling to promote tolerance to ischemic brain injury. J Neurosci 31: 8456–8463

Stevens SL, Vartanian KB, Stenzel-Poore MP (2014) Reprogramming the response to stroke by preconditioning. Stroke 45: 2527–2531

Stoll G, Schroeter M, Jander S, Siebert H, Wollrath A, Kleinschnitz C, Bruck W (2004) Lesion-associated expression of transforming growth factor-beta-2 in the rat nervous system: evidence for down-regulating the phagocytic activity of microglia and macrophages. Brain Pathol 14: 51–58

Sun X, Lindsay J, Monsein LH, Hill PC, Corso PJ (2012) Silent brain injury after cardiac surgery: a review: cognitive dysfunction and magnetic resonance imaging diffusion-weighted imaging findings. J Am Coll Cardiol 60: 791–797

Supek F, Bosnjak M, Skunca N, Smuc T (2011) REVIGO summarizes and visualizes long lists of gene ontology terms. PLoS One 6: e21800

Tang SC, Arumugam TV, Xu X, Cheng A, Mughal MR, Jo DG, Lathia JD, Siler DA, Chigurupati S, Ouyang X et al (2007) Pivotal role for neuronal Toll-like receptors in ischemic brain injury and functional deficits. Proc Natl Acad Sci U S A 104: 13798–13803

Tasaki K, Ruetzler CA, Ohtsuki T, Martin D, Nawashiro H, Hallenbeck JM (1997) Lipopolysaccharide pre-treatment induces resistance against subsequent focal cerebral ischemic damage in spontaneously hypertensive rats. Brain Res 748: 267–270

Team RC (2019) R: A language and environment for statistical computing. R Foundation for Statistical Computing

Thomalla G, Simonsen CZ, Boutitie F, Andersen G, Berthezene Y, Cheng B, Cheripelli B, Cho TH, Fazekas F, Fiehler J et al (2018) MRI-Guided Thrombolysis for Stroke with Unknown Time of Onset. N Engl J Med 379: 611–622

Vanheule V, Metzemaekers M, Janssens R, Struyf S, Proost P (2018) How post-translational modifications influence the biological activity of chemokines. Cytokine 109: 29–51

Veldhuis WB, Derksen JW, Floris S, Van Der Meide PH, De Vries HE, Schepers J, Vos IM, Dijkstra CD, Kappelle LJ, Nicolay K et al (2003a) Interferon-beta blocks infiltration of inflammatory cells and reduces infarct volume after ischemic stroke in the rat. J Cereb Blood Flow Metab 23: 1029–1039

Veldhuis WB, Floris S, van der Meide PH, Vos IM, de Vries HE, Dijkstra CD, Bar PR, Nicolay K (2003b) Interferon-beta prevents cytokine-induced neutrophil infiltration and attenuates blood-brain barrier disruption. J Cereb Blood Flow Metab 23: 1060–1069

Virani SS, Alonso A, Aparicio HJ, Benjamin EJ, Bittencourt MS, Callaway CW, Carson AP, Chamberlain AM, Cheng S, Delling FN et al (2021) Heart Disease and Stroke Statistics-2021 Update: A Report From the American Heart Association. Circulation 143: e254–e743

Werner Y, Mass E, Ashok Kumar P, Ulas T, Handler K, Horne A, Klee K, Lupp A, Schutz D, Saaber F et al (2020) Cxcr4 distinguishes HSC-derived monocytes from microglia and reveals monocyte immune responses to experimental stroke. Nat Neurosci 23: 351–362

Zhu G, Wang X, Chen L, Lenahan C, Fu Z, Fang Y, Yu W (2022) Crosstalk Between the Oxidative Stress and Glia Cells After Stroke: From Mechanism to Therapies. Front Immunol 13: 85241

